# Recurrent circuit based neural population codes for stimulus representation and inference

**DOI:** 10.1101/2020.11.18.389197

**Authors:** Wen-Hao Zhang, Si Wu, Krešimir Josić, Brent Doiron

## Abstract

A large part of the synaptic input received by cortical neurons comes from local cortico-cortical connectivity. Despite their abundance, the role of local recurrent connections in cortical function is unclear, and in simple coding schemes it is often the case that a circuit with no recurrent connections performs optimally. We consider a recurrent excitatory-inhibitory circuit model of a cortical hypercolumn which performs sampling-based Bayesian inference to infer latent hierarchical stimulus features. We show that local recurrent connections can store an internal model of the correlations between stimulus features that are present in the external world. When the resulting recurrent input is combined with feedforward input it produces a population code from which the posterior over the stimulus features can be linearly read out. Internal Poisson spiking variability provides the proper fluctuations for the population to sample stimulus features, yet the resultant population variability is aligned along the stimulus feature direction, producing *differential correlations*. Importantly, the amplitude of these internally generated differential correlations is determined by the associative prior in the model stored in the recurrent connections. This provides experimentally testable predictions for how population connectivity and response variability are related to the structure of latent external stimuli.

## Introduction

In an uncertain and changing world, it is imperative for the brain to reliably represent and interpret stimuli. A wrong decision based on a faulty stimulus representation could endanger an animal’s life. The sensory cortex is essential for the representation of the sensory world^1;2^, with individual neurons tuned to stimulus features so that the population as a whole can code for a richly structured sensory scene. However, two central cortical circuit features remain to be properly integrated into a coherent representation framework. First, neuronal responses in sensory cortices are often noisy, with the same stimulus evoking starkly different responses across trials ^3;4^. If such response variability is unrelated to sensory stimuli, it would be detrimental to their representation in cortical circuits ^2^. Why cortical responses are so variable when isolated cortical neurons can respond reliably remains a mystery^5^. Second, the primary source of synaptic inputs to cortical neurons is not from upstream centers which presumably carry sensory signals, but rather from recurrent pathways between neurons in the same hierarchical layer ^6–8^. While such recurrent connections are organized about a stimulus feature axis ^9;10^, it is not obvious whether and how their presence improves overall representation. Here we propose a biologically motivated inference coding scheme where these two cortical circuit features are essential for optimal stimulus representation.

Numerous studies have shown that cortical sensory processing can be understood in terms of Bayesian inference^11–17^. In such models, sensory cortex infers and synthesizes a description of the possible stimuli that could have given rise to the sensory inputs, rather than generating a single stimulus estimate. To perform Bayesian inference, the sensory cortex needs to store a model describing how external stimuli and the resulting sensory inputs are generated. Once a sensory input is received, cortical dynamics inverts this internal model in a process termed ‘analysis-by-synthesis’ ^13^. The neural response represents a posterior distribution over the input stimulus and unobserved (latent) variables^16;17^. We propose that recurrent (local) connections in cortical circuits represent such an internal model ^18^, and that internally generated variability is a source of fluctuations needed for generating samples from the posterior distribution, ^17;19;20^. The challenge in this approach is to determine how the structure of connectivity and population variability should be set so as to accurately represent the posterior distribution.

To tackle this challenge we consider a hierarchical generative model ^13^, where sensory inputs are generated from a local stimulus feature and a global, latent context feature. We assume that the local and global aspects of a scene are correlated, so that knowledge of one provides information about the other. Examples would be the direction of a breeze (global) and a leaf’s movement in that breeze (local), or the orientation of a tree trunk (global) and orientation of bark texture on a section of the trunk (local). We propose a recurrent network model that interprets these inputs by implementing sampling-based inference over the stimulus and context features. The inferred posterior over the latent features is represented in a subspace of the neuronal population response, and can be linearly read out by a downstream decoder. Importantly, we find that the recurrent connections store the associative prior between stimulus and context features, and that internal Poissonian variability can drive sampling in the feature subspace. The best approximation of the posterior over the stimulus and context features is achieved in a network with recurrent connections whose strength increases with the degree of correlation between context and local sensory inputs. In other words, a strongly coupled network is needed for optimal inference in a well-structured world, whereas an unconnected network would work best in a world without structure.

We provide testable experimental predictions about how the brain represents the structure of the world to interpret sensory information. We propose that a neural signature of network sampling is an internally generated response covariability along the direction of stimulus features, known as differential correlations ^2;21^. The amplitude of these internally generated differential correlations is determined by the associative prior between stimulus and context features in the internal model stored in the recurrent connections. Thus measurable differential noise correlations can be used as a proxy for strong recurrent connections organized along the stimulus tuning dimension. These correlations can be controlled by adjusting the structure of the external world.

## Results

### Recurrent circuitry does not improve conventional neural codes

Neural coding is the process by which a stimulus feature, *s*, is encoded in neuronal activity, **r**, which is subsequently decoded to provide an estimate of the feature, *ŝ* (Fig. 1a, top) ^22^. In this view, it is reasonable to expect that neuronal circuitry is adapted to provide an accurate feature estimate. However, we next show that in simple coding schemes recurrent connectivity at best does not affect stimulus estimation, and at worst degrades such estimates, raising the question of its purpose.

**Figure 1:**
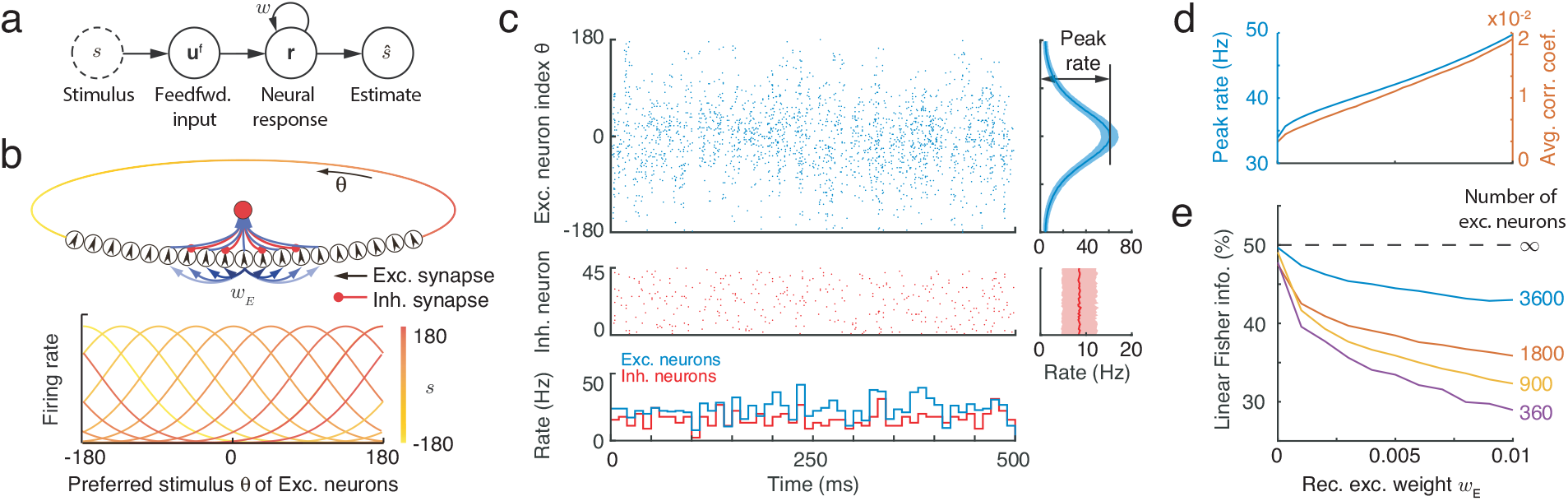
A network with structured recurrent connections limits linear Fisher Information (LFI) about external stimuli. (a) A schematic diagram showing how a stimulus, *s*, is encoded in a neuronal response, **r**. A stimulus estimate *ŝ* can be decoded from **r**. (b) A recurrent ring model (top) with population activity of excitatory neurons, **r**, dependent on a stimulus, *s* (bottom). Information about the stimulus can be obtained from the activity of E neurons. (c) Example realization of population spiking activity of excitatory and inhibitory neurons in the network. Right: the firing rate of neurons averaged over time, with shaded region encompassing one standard deviation from the mean. (d) The population responses measured in peak firing rate (left axis) and spike count correlations averaged over excitatory neurons (right axis). (e) For finite size networks (colored lines; ratio of excitatory to inhibitory neurons kept constant) the LFI decreases as *w_E_* increases. In the limit of infinite network size LFI does not depend on *w_E_* (dashed line).

In classical models, stimulus encoding is performed by a neuronal population, with individual neurons tuned to a preferred stimulus value. The preferred values across the population cover a range of potential stimuli ^22–24^(Fig. 1b, bottom). If the stimulus, *s*, ranges over a periodic domain (such as the orientation of a bar in a visual scene, or the direction of an arm reach) then it is commonly assumed that the neurons’ preferred stimuli are distributed on a ring (Fig. 1b, top). Neurons with similar feature preferences tend to be coupled more strongly ^9;10^, so that network connectivity aligns with the ring topology (Fig. 1b, top, arrows). We implemented such a model with structured connectivity between excitatory (E) neurons, and unstructured inhibition (I) that acts only to stabilize network activity ^25^. Hence, only E neurons in the network are tuned to a stimulus feature (Fig. 1b, bottom). Further, for simplicity we assumed a linear (or linearized) neuronal transfer and synaptic interactions (Eqs. 19–20), and Poisson spike responses (Eqs. 31–34). We presented the network with a stimulus with feature *s* = 0, resulting in elevated activity of E neurons with the corresponding preference (Fig. 1c). As expected, an increase in the strength of recurrent excitatory connections increases both the firing rates and the trial-to-trial pairwise covariability (i.e. noise correlations) in the responses ^3^ (Fig. 1d). This canonical, mechanistic network model has been widely used to explore cortical network dynamics and coding ^25–27^.

We considered the impact of recurrent connectivity on the accuracy of the decoded feature estimate, *ŝ*, using a conventional coding framework. Towards this end, we measured the linear Fisher Information (LFI) of *s* in the population response of E neurons (see details in Eq. S24 in Supplementary Information). The inverse of the LFI provides a lower bound on the squared error between *s* and *ŝ* made by a linear decoder^1;2;21–23;28^, and it is closely related to an animal’s performance in discrimination tasks widely studied in experiments^2;22^. We found that for networks with large numbers of decoded neurons LFI is not affected by the peak recurrent strength *w_E_* (Fig. 1e, dashed line), and converges to the information about *s* provided by the feedforward input ^21^. For the simple network and stimulus world we consider (Fig. 1a,b) this result can be readily understood: Since in our network a neuron’s mean response is a linear function of its feedforward and recurrent inputs, the entire network acts as a linear filter of the stimulus *s*. An (invertible) linear filter can neither augment nor degrade the stimulus information available to be decoded by a linear readout (as LFI measures; see Eq. S15 in Supplementary Information). Thus, LFI does not depend on the details of the filter, or in this case on the strength of recurrent coupling *w_E_*. However, networks with a finite number of neurons produce internal noise which can degrade information about the stimulus, and hence reduce the LFI (Fig. 1e, colored lines) ^21;28^. For such finite networks this noise is shared between neurons and thus an increase with coupling strength, *w_E_*, in turn reducing LFI. In sum, recurrent connectivity does not improve stimulus representation in terms of increasing LFI. Since synaptic coupling is biologically expensive, a network that most accurately and cheaply represents stimulus information is one that does not have any recurrent connections (i.e., *w_E_* = 0).

What is then the purpose of the extensive recurrent connections between neurons^6–8;10^? There are at least three possibilities. First, while recurrent connectivity does degrade stimulus representation, it might underlie other stimulus processing functions essential for cortical computations. Second, LFI and the linear network interactions we assumed are overly simplistic, and nonlinear network interactions^28^ or nonlinear decoders ^29^ may be required to properly model population coding, and the performance of neural codes may be optimized at nonzero coupling strengths. The final possibility is that it is the stimulus world we used to probe neural coding (Fig. 1a) that is overly simplistic, and recurrent cortical coupling in our linear network can improve cortical codes that represent more complex stimulus worlds. We next develop an extended framework that shows that the last possibility is plausible, but does not exclude the first two.

### A hierarchical framework of information processing in recurrent circuit

Classical population codes provide a point estimate (i.e., a single value) of a latent stimulus, *s*, transmitted via a feedforward sensory pathway (Fig. 1a). However, numerous studies suggest that the brain is performing Bayesian inference^11–13;15;32;33^, and thus synthesizes and estimates the probability of a range of possible latent stimuli. To accomplish this task neural circuits must approximate a posterior distribution over the latent stimulus from the sensory input, **u**^f^, and any stored prior information. Such computations can be complex, and require more biological machinery than point estimation. Here we propose a circuit based implementation of Bayesian inference, paying particular attention to the role of recurrent wiring.

In order for the circuit computation to be non-trivial, the stimulus has to come from a world with sufficiently rich structure. We therefore consider a hierarchical generative model (Fig. 2a) that describes how a feedforward input, **u**^f^, received by the cortical circuit is generated from stimulus and context features in the outside world. We assume that the first stage of our model consists of a stimulus feature, *s*, and a context feature, *z*, both of which are one dimensional for simplicity. The structure of the world is described by the joint distribution, *p*(*s, z*). In the example of a V1 hypercolumn the stimulus feature, *s*, could be the orientation of the visual texture within a classical receptive field (local information) of a hypercolumn of V1 neurons, while the orientation within a non-classical receptive field of these cells could describe the corresponding context feature, *z* (Fig. 2a). We assume that the conditional distribution of the stimulus feature, for a given context, 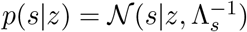, is Gaussian with precision (inverse of the variance) equal to Λ_*s*_. For simplicity, we assume that the prior distribution over the context feature, *p*(*z*), is uniform (Fig. 2c). As we show in the Discussion, this assumption is not essential for our main conclusions. While the uniform prior over the context feature, *z*, implies that the marginal prior over *s*, is also uniform, the joint prior over stimulus and context features, *p*(*s, z*), can have non-trivial structure. Specifically, if the two features are positively correlated, the joint prior density is concentrated around the diagonal (Fig. 2c). The width of the band is determined by the precision Λ_*s*_, which measures how strongly the context, *z*, and the stimulus feature, *s*, are related.

**Figure 2:**
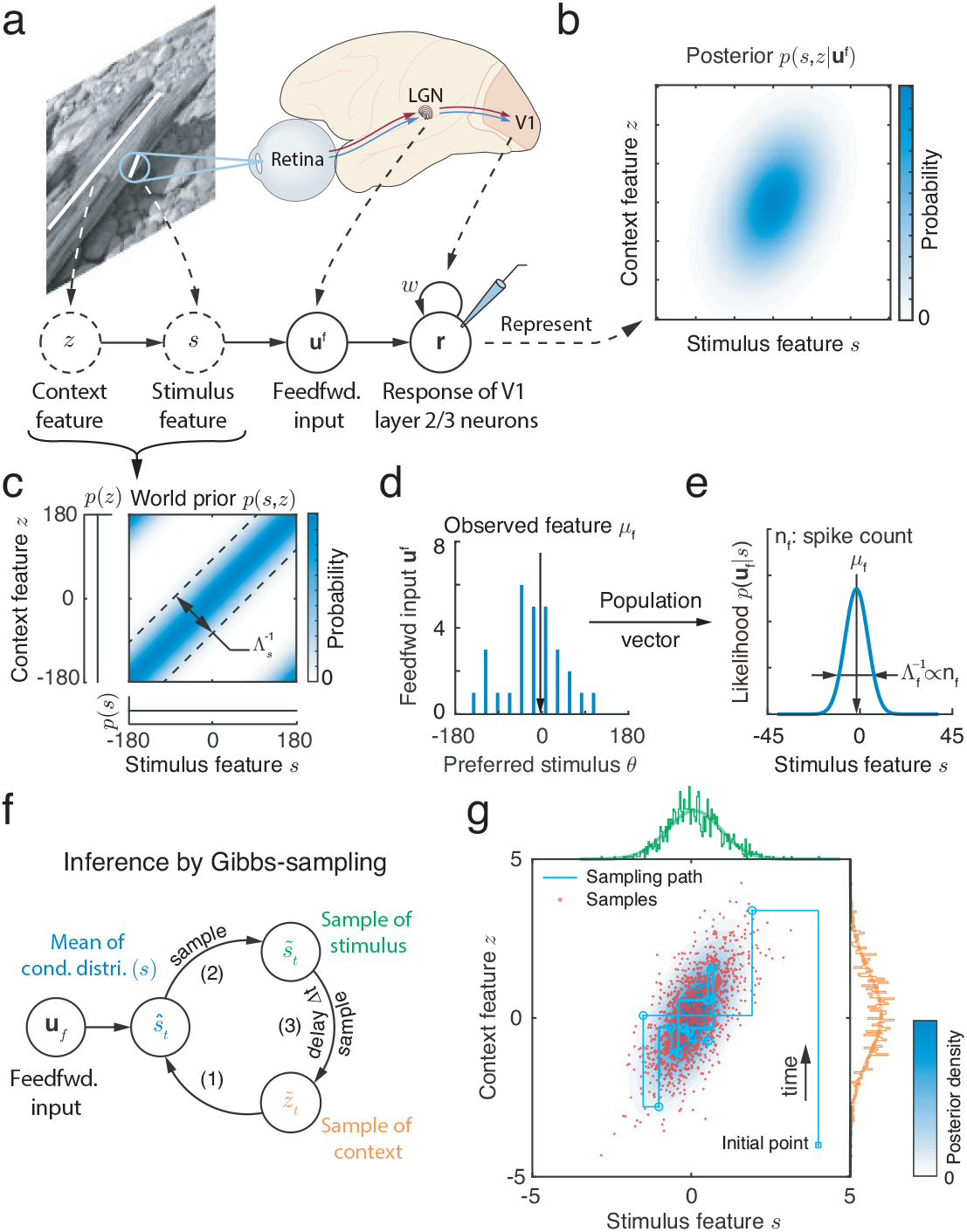
A hierarchical generative model and inference by Gibbs sampling. (a) An example of sensory feedforward input generation: The context feature, *z,* is the orientation of the tree trunk, and the stimulus feature, *s*, is the orientation of the bark texture located in the classical receptive field of a V1 hypercolumn. Feedforward input, **u**^f^, conveys stimulus information to V1 neurons. The evoked response, **r**, represents the joint posterior over stimulus and context, *p*(*s, z*~**u**^f^). Solid circles: observations and responses in the brain; dashed circles: latent variables in the world. (Natural image and brain schematic adapted from 30 and 31 respectively). (b) The joint posterior *p*(*s, z*~**u**^f^) over stimulus and context can be represented by the neuronal response, **r**. (c) The joint prior over the global context, *z*, and local stimulus, *s*, has more mass around the diagonal. The width of this band is controlled by Λ_*s*_, and determined by the correlation between context and stimulus. (d) A realization of feedforward spiking inputs, **u**^f^, defined as independent Poisson random variables with intensity dependent on stimulus, *s*. (e) The parameters of the likelihood function of *s* can be linearly read out from **u**^f^ using a population vector (Eq. 12). (f) The posterior over context and stimulus can be approximated via Gibbs sampling (Eqs. 2a–2c) by iteratively generating samples of *s* and *z* from their respective conditional distributions. (g) The resulting approximations of the joint and marginal posterior over the latent stimulus feature, *s*, and context feature, *z*. Light blue contour: the posterior distribution (Eq. 14); Red dots: Samples obtained using Gibbs sampling.

The second stage of the model describes stimulus transmission via the primary sensory pathway (from *s* to **u**^f^ in Fig. 2a). The feedforward input, **u**^f^, is represented by a vector of independent Poisson spike counts (Eq. 10) that are received by *N_E_* excitatory (E) neurons in the recurrent circuit. The mean intensity of each component of **u**^f^ has a Gaussian tuning profile over the stimulus feature, *s*, so that 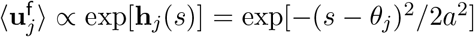. We assume that the preferred stimuli of the feedforward inputs, 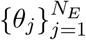, are uniformly distributed over the stimulus feature range, and that the widths of all tuning curves equal a constant, *a* ^22;34;35^. We refer to the subspace in the space of sensory inputs spanned by the vector (*θ*_1_*, θ*_2_,…,*θ_N_E*) as the *stimulus feature subspace*. The Gaussian tuning of the input is consistent with the broad, bell-shaped orientation tuning in V1 (Fig. 1b, bottom), and conditional independence of the Poisson input is consistent with the observation that LGN and V1 simple cells remove redundancy in the input ^30^.

Combining these two input stages provides a complete description of the feedforward input, **u**^f^, (see details in Eq. 11),

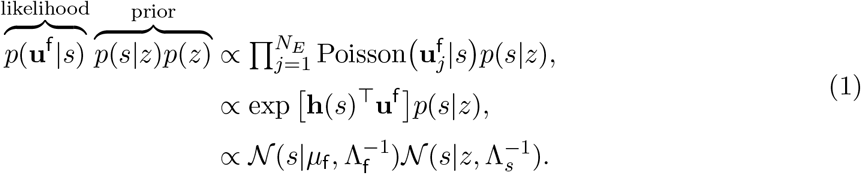

The input vector, **u**^f^, represents a Gaussian likelihood over the stimulus feature, *s*^34;35^, i.e., 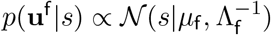. This is a consequence of the Gaussian tuning, and the assumption that the preferred stimuli of feedforward inputs are uniformly distributed. The likelihood of the stimulus, *s*, is thus determined by the mean, *μ*_f_, and precision, Λ_f_, both of which can be read out linearly from the input, **u**^f^, using a population vector ^24;34^ (Eq. 12): The position (center of mass) of **u**^f^ in the stimulus feature subspace (x-axis in Fig. 2e) represents the mean, *μ*_f_, which corresponds to a noisy observation of stimulus feature (Fig. 2d-e). Further, the precision, Λ_f_, is proportional to the sum of input spike counts, 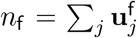 (Eq. 12). We next ask how a recurrent neural circuit can approximate the joint posterior distribution over the stimulus features *s* and *z* from sensory input, **u**^f^ (Fig. 2b).

### Sampling based inference of the joint stimulus posterior

Given the generative model with the additional assumptions captured in Eq. (1), the joint posterior over stimulus and context features, *p*(*s, z*|**u**^f^), is a multivariate normal distribution (See Eq. 14). In Bayesian theories of perception, the computed approximation of this posterior captures an observer’s perception of stimulus and context ^11;12;14;16;32;33^. We next show how recurrent circuit dynamics can be used to approximately compute, and represent this posterior distribution ^1;16;17^.

We propose that Gibbs sampling, a stochastic algorithm which generates samples from a probability distribution ^36^, can be used to approximate the posterior over the features *s* and *z* given sensory input, **u**^f^. This algorithm requires a mechanism for generating random numbers, and is thus well suited for implementation in cortical circuits given the prominent stochasticity of neuronal activity in cortex ^13;17;19;37–40^. Gibbs sampling starts with an initial guess for the value of the two latent features, and proceeds by alternately fixing a sampled value of one feature, and updating the other by drawing a sample from the respective conditional distribution. More precisely, to approximate the posterior distributions over *s* and *z*, Gibbs sampling proceeds by generating a sequence of samples, 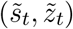 indexed by time *t*. We consider discretized time with *t* = Δ*t*, 2Δ*t*,…,*T*, where Δ*t* is the time increment between successive samples, *T* is the sampling duration, and thus *T/*Δ*t* are the number of samples. This discrete construction merely simplifies exposition; extensions to continuous time sampling are straightforward. Given the generative structure of the stimulus described in the previous section, such sampling can be implemented through a recursive iteration of the following algorithm (Fig. 2f and Eq. (15)),

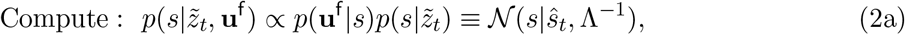

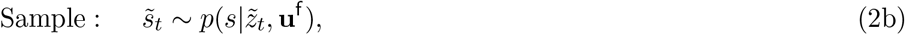

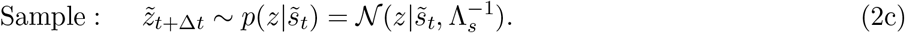

The samples are generated by alternately fixing the values of the two variables, so that sampling trajectories alternate between horizontal and vertical jumps in parameter space (cyan line in Fig. 2g). The set of sampled values (red dots in Fig. 2g) defines a distribution, 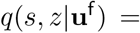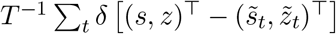, where *δ*(·) is the Dirac delta function and T denotes vector transpose. The distribution *q*(*s, z*|**u**^f^) approximates the true posterior distributions *p*(*s, z*|**u**^f^) (blue contour map in Fig. 2g, Eq. 14) ^36^. Moreover, to approximate the marginal posterior distribution over one of the features, say *p*(*s*|**u**^f^), we can keep only samples 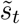 to obtain an approximating distribution *q*(*s*|**u**^f^) (compare the two green lines at the margin in Fig. 2g).

### Bayesian inference in a recurrent network of spiking neurons

If the brain performs Bayesian inference it is important to understand how the requisite probabilistic computations are implemented in cortical circuits. We next show how the proposed Gibbs algorithm for alternate sampling of the stimulus feature, *s*, and the context feature, *z* can be naturally implemented in a recurrent network with internally generated variability. We first explain the mechanism using a linear network model consisting of only E neurons (Fig. 3a, Eqs. 19–20), and later demonstrate how it can be implemented in a network consisting of both E and I neurons (Fig. 1). The response of the E population at time *t*, **r**_*t*_, is a vector of spike counts, with each entry sampled from a Poisson distribution with parameters given by the vector of firing rates ***λ**_t_*. We assume that these firing rates are a sum of the sensory feedforward input, **u**^f^ (Eq. 1), and recurrent inputs 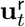.

**Figure 3:**
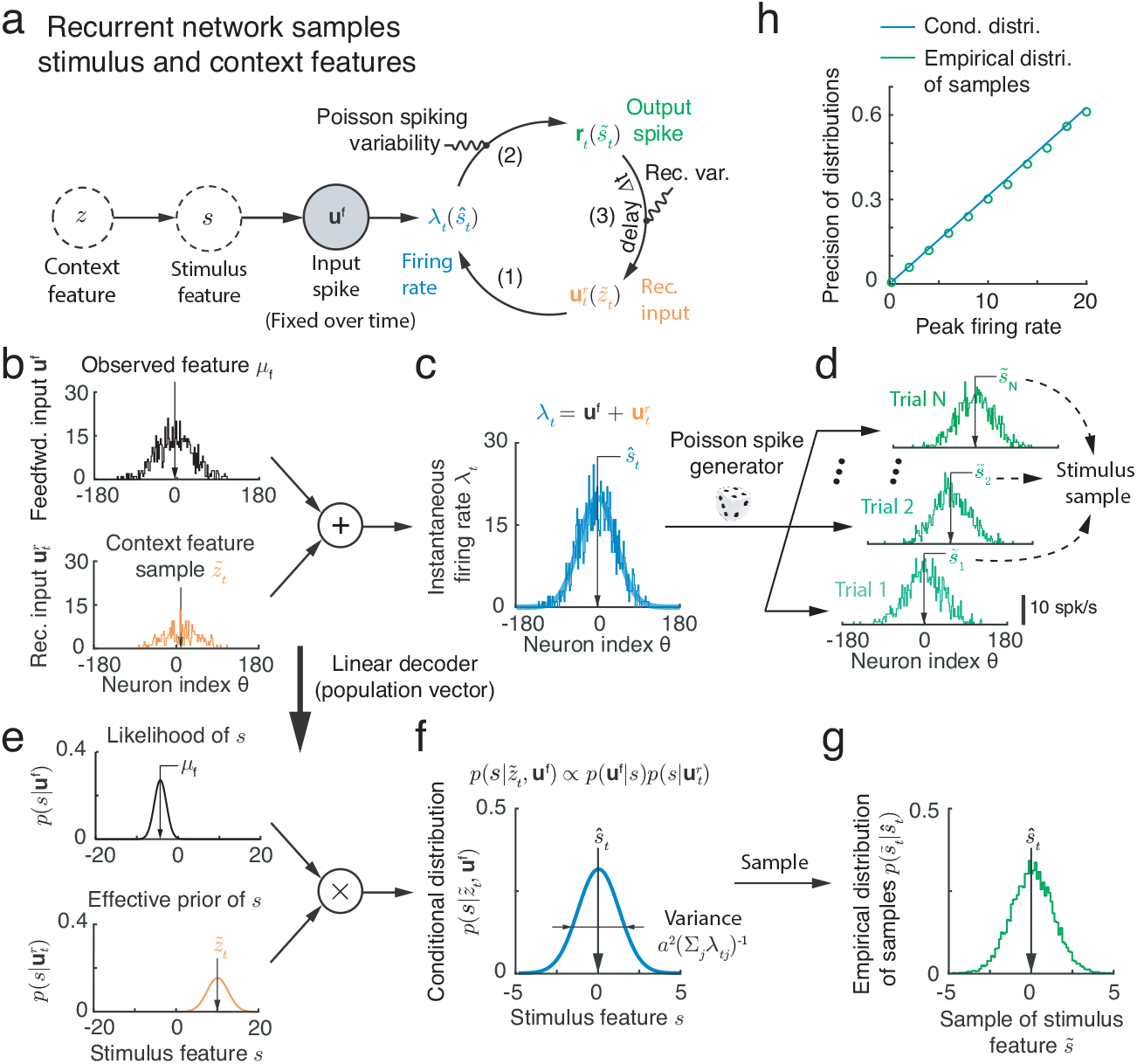
A recurrent circuit with internal Poisson variability can implement latent feature sampling. (a) Schematic of recurrent circuit dynamics, in which stimulus, *s*, and context, *z*, features are encoded respectively by the population response, **r**, and recurrent inputs **u**^r^. Poissonian spiking and recurrent variability produce the fluctuations needed for sampling stimulus and context features, respectively. (b-d) The sum of feedforward and recurrent inputs (b) yields the instantaneous firing rate (c), which determines the mean of independent Poisson spike counts (d). (e-g) The distributions and samples of stimulus features are linearly read out from corresponding population inputs and responses shown in (b-d). The empirical distribution of samples (g) from Poisson spike generation in (d) is consistent with the conditional distribution (f) represented by instantaneous firing rate (c). (h) The peak firing rate determines the precision of the conditional distribution of stimulus feature (f) and the variability of stimulus feature samples.

We next need to resolve several fundamental questions: How does the recurrent circuit represent the generative model that needs to be inverted to infer the latent variables (Eq. 1)? How are the conditional distributions, and samples from the posterior over the features (Eqs. 2a–2c) represented in neural circuitry and activity? Random sampling requires internally generated variability whose statistics should be determined by conditional distributions that change with each sample (Eq. 2b–2c). Is there a universal and robust circuit mechanism to generate such variability? We address these questions next.

### Representing the posterior in the recurrent network

We first show how a recurrent network can store an internal representation of the generative model by comparing Gibbs sampling (Fig. 2f) to recurrent network dynamics (Fig. 3a): A sample of the stimulus feature, 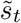, will be represented by the response of E neurons in the recurrent network, **r**_*t*_, while a sample of the context feature, 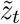, will be represented by the recurrent inputs, 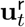. In the network the sum of feedforward and recurrent inputs (Fig. 3, step 1) is used to compute the conditional distribution of the stimulus feature (Eq. 2a). For this to hold, we make two further assumptions: First, we assume the mean recurrent input, 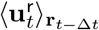, averaged over realizations (trials) conditioned on the response **r**_*t*−Δ*t*_, has the same tuning profile as the feedforward inputs (Eq. 1) but centered at the stimulus feature sample 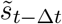 represented by **r**_*t*−Δ*t*_, i.e., 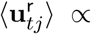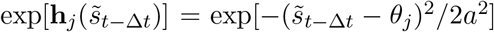. Second, given the response **r**_*t*−Δ*t*_, we assume that the recurrent inputs, are independent, and have Poisson-like variability (the variance equals the mean, see Eq. 21). Under these assumptions, the stimulus feature, *s*, given recurrent input 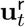, follows a normal distribution,

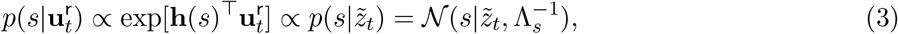

whose mean, 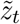, and precision, Λ_*s*_, can be read out linearly from 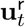 (Fig. 3b and e, bottom-left; Eq. 23). Since the recurrent input encodes a sampled value of context feature, we refer the subspace in the space of recurrent inputs spanned by the vector 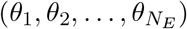 as the *context feature subspace*. By construction, the recurrent and feedforward inputs represent conditional distributions of the stimulus feature, *s*, with the same basis function, **h**(*s*) (logarithm of the tuning curve). Therefore summing the feedforward and recurrent inputs to a neuron corresponds to multiplying the likelihood, *p*(**u**^f^ |*s*), and the conditional distribution 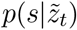 (Eq. 3; Fig. 3c and f),

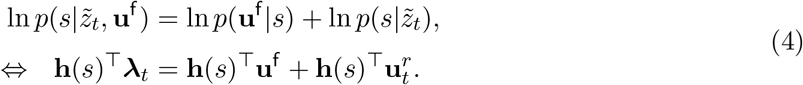

The firing rate of the neurons in the recurrent circuit, ***λ****_t_*, thus represents the instantaneous conditional distribution, 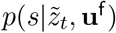, and has the same basis function, **h**(*s*), as that of the feedforward and recurrent inputs (Eq. 2a). Table 2 summarizes how inputs and instantaneous firing rate represent the distributions over the stimulus features.

### Internal Poisson variability drives sampling in stimulus feature subspace

A neural circuit that implements sampling must generate specific level of variability in the feature subspace (see Eq. 2b–2c). A sample from the conditional stimulus distribution (Eq. 2b) can be written as 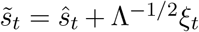, where *ŝ*_*t*_ and Λ are the mean and precision of the conditional distribution 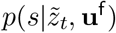 encoded in the mean firing rates, ***λ**_t_* (Eqs. 2a and 4), and *ξ_t_* are independent samples from a standard normal distribution. If spiking responses, **r**_*t*_, represent stimulus samples, 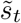, then the recurrent network must generate neuronal responses whose projection onto the stimulus feature subspace follows a Gaussian distribution with variance Λ^−1^.

Poisson-like spiking is observed across cortex ^4;22^, and is thus a natural candidate for the requisite variability. We show that when the instantaneous firing rate, ****λ***_t_*, represents the conditional distribution, 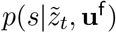 in Eq. (4), then independent Poisson spike generation produces samples with the correct statistics in feature subspace (Fig. 3d and g, details in Methods),

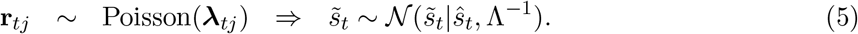

A sample from the conditional stimulus feature distribution, 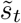, can be read out from the population response, **r**_*t*_, using a linear decoder, 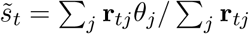. The posterior mean, *ŝ_t_*, is related to the mean firing rates, ***λ**_t_*, in the same way. The weights in this decoder are determined by the basis functions (logarithm of tuning), **h**(*s*), as in Eq. (4). Importantly, the variability of stimulus feature samples, Λ^−1^, is determined by the sum of the instantaneous firing rates, Σ_*j*_**λ**_*tj*_.

To confirm that Poisson spike generation leads to sampling from the correct stimulus feature distribution, we fixed a realization of instantaneous firing rates, ***λ****_t_* (Fig. 3c), which corresponds to fixing the conditional distribution of the stimulus feature (Eq. 4), and then generate independent Poisson spikes over trials (Fig. 3d). In each trial, a sample of 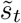 was read out by using a population vector (Fig. 3d, Eq. 5). The empirical distribution of the samples, 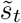, collected over trials (Fig. 3g) matched the conditional distribution represented by the instantaneous firing rate, ***λ****_t_* (Fig. 3c and f, Eq. 4). We have thus shown that internally generated Poisson variability can provide a source of fluctuations for sampling from the the conditional distribution over the stimulus feature. Further, we confirm that the overall firing rate of the neural population determines the variability of the stimulus feature samples (Fig. 3h). An advantage of this implementation is that the sampling variability, Λ^−1^, is determined by firing rates, and can thus change over short times scales.

### Recurrent connections represent the relation between stimulus and context features

To generate samples from the correct joint distribution of stimulus and context features, the mean and variance of the recurrent input, **u**^r^, to neurons in the circuit must satisfy Eq. (3). First, the input, **u**^r^, and the population response, **r**, should both have Gaussian tuning with the same tuning width over the stimulus feature, *s* (see Eqs. 21 and 24). Second, the mean and variance of the recurrent input conditioned on the population responses, **r**_*t*_, need to be equal.

We show that the first requirement is satisfied when the weights between neurons in the recurrent circuit are self-connected with homogeneous connection strength (see Eq. 26). The optimal recurrent weight to correctly sample the posterior is

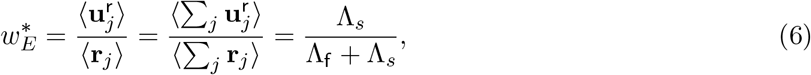

where Λ_*s*_ is the precision of the stimulus prior and Λ_f_ is the precision of the feedforward input distribution *p*(**u**^f^ |*s*). The recurrent weight, 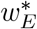, thus encodes the relation between the stimulus feature, *s*, and the context feature, *z*. An increase in correlation between *s* and *z* (reflected by a narrower diagonal band in Fig. 2c) requires an increase in the recurrent weight 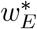 to satisfy Eq. (3). However, when context and stimulus are uncorrelated (Λ_*s*_ = 0), then recurrent interactions are not needed (i.e.,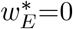). This is in agreement with the example we chose to motivate our approach, where LFI was maximized in the absence of recurrent connectivity (Fig. 1e).

To fulfill the second requirement, the recurrent interactions in the network need to exhibit Poisson-like variability (Fig. 3a, step 3; Eq. 19), which could be the result of noisy synaptic transmission^41;42^. The resulting recurrent input, 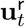, introduces the variability needed to generate samples from the conditional context feature (*z*) distribution (compare step 3 in Fig. 2d to the recurrent interactions in Fig. 3a). This sampling mechanism is equivalent to the one we proposed for generating samples from the stimulus feature distribution (Eq. 5)

### A stochastic E-I network implements sampling in feature subspace

To confirm that a biologically realistic neural circuit can carry out Gibbs sampling, we implemented the proposed algorithm in a recurrent network consisting of E and I Poisson neurons with synaptic interactions (See details in Eqs. 31–34). The feedforward and recurrent circuitry of this network was identical to the one used to probe the effect of recurrent coupling on a linear estimation of stimulus *s* (Fig. 1). We assumed that the response properties of the individual neurons satisfy the representation requirements we described previously: The tuning of the recurrent inputs, **u**^r^, and neuronal responses, **r**, followed the same Gaussian profile as the feedforward input, **u**^f^ (Eq. 4). The recurrent inputs in the network exhibited independent Poisson-like variability given a spiking response (Eq. 32). Thus, the model included three sources of variability (Fig. 4a): The external variability in feedforward inputs **u**^f^ which determines the likelihood function of stimulus feature (Eq. 1), internal variability due to spike generation driving sampling from the stimulus feature distribution, and variability in the recurrent interactions driving the sampling from the context feature distribution. We next demonstrate that with these sources of variability the network is able to infer stimulus and context features.

**Figure 4:**
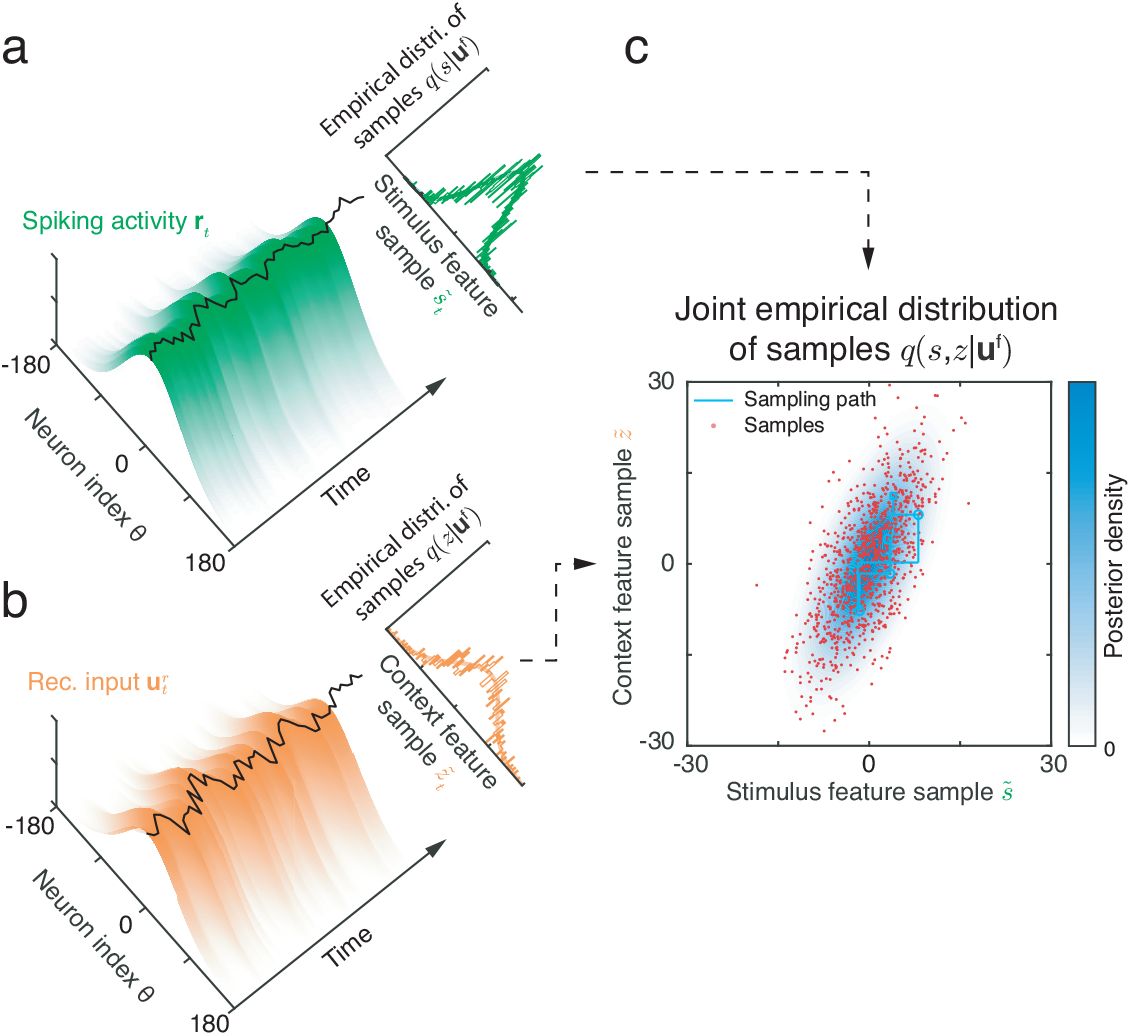
Reading out the posterior over latent features from spiking activity and recurrent inputs in a model cortical circuit. (a-b) Spiking activity, **r**_*t*_ (a), and recurrent input 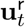 (b) evolve stochastically. Their projection onto the feature subspace (black curve overlaid on the top of the surface), can be read out linearly from the population activity and interpreted as a sample from the joint posterior distribution over stimulus and context features (Eqs. 2b-2c). Top right insets: the empirical marginal distributions of samples and marginal posteriors of features (smooth lines). (c) The joint value (red dots) of instantaneous samples of stimulus feature (black curve on the surface in a), and context feature (black curve on surface in b) represent samples from the joint posterior of the stimulus and context. The true joint posterior distribution is shown in blue.

### Population responses of a model network generates feature samples

To show that the model recurrent network produces samples from the joint posterior over stimulus and context features, we simulated its response to a fixed feedforward input **u**^f^ generated according to Eq. (1). Response variability was due to the combination of Poissonian spiking and noisy recurrent interactions. Fixing **u**^f^ separates external and internal noise sources in the network, allowing us to directly compare samples generated by the network with the posterior conditioned on a fixed input, **u**^f^ (Eq. 14; Fig. 1b). Since inhibition is unstructured (see Methods) only E neurons are selective for the stimulus and context features in the E-I network (Fig. 1c), implying that we only need to read out the samples of features from E neurons. To simplify notation, we therefore suppress subscripts denoting neuronal type in the main text, and denote responses and recurrent inputs to E neurons by **r**_*t*_ and 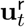, respectively.

By construction, samples of stimulus and context features are represented respectively by the position of the instantaneous responses, **r**_*t*_, and the recurrent inputs, 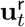, in feature subspace (black curves overlaid on the top of green and orange population responses in Fig. 4a and b). Hence the samples of stimulus and context features can be read out separately from **r**_*t*_ and 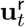 using a population vector (Eqs. 5 and 23; Fig. 3b-g). The resulting trajectory of samples in the direct product of the stimulus and context feature subspace exhibits the characteristics of Gibbs sampling (Fig. 4c, cyan line) as the spiking activity and recurrent inputs are updated iteratively in the network. The empirical distribution produced by the collection of samples, 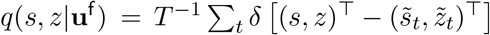, (Fig. 4c, red dots) provides a good approximation of the joint posterior over stimulus and context features (Fig. 4c, blue contour).

### Optimal inference by the network with the correct recurrent weight

To produce correct samples from the joint posterior over stimulus and context features, the recurrent connectivity must support network activity which correctly captures the generative model of the stimulus (Eq. 6). We next verified that setting the recurrent weights to an optimal value allows for sampling from the correct posterior distribution, and that these optimal weights are determined by the prior precision, Λ_*s*_, as suggested by the theory we developed earlier (Eq. 6).

Towards this end, we fixed all parameters in the generative model (Eq. 1) and changed only recurrent weights in the model network. For simplicity, we only varied the peak excitatory strength, *w_E_* (Eq. 33), and maintained stable activity by fixing the ratio of excitatory and inhibitory synaptic weights. The joint distributions of samples produced by the recurrent network varied considerably as we varied the peak recurrent weights, *w_E_* (Fig. 5a). Changing *w_E_* did not change the mean, but did affect the variance and covariance of the distribution of samples, and hence the inferred distribution over the features (Fig. 5a). Stronger weights, *w_E_*, increased the variance in the marginal distribution over stimulus features, while the variance over context feature samples decreased. Moreover, the orientation of the joint distribution rotated clockwise, increasing the correlations between the samples 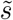 and 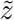.

**Figure 5:**
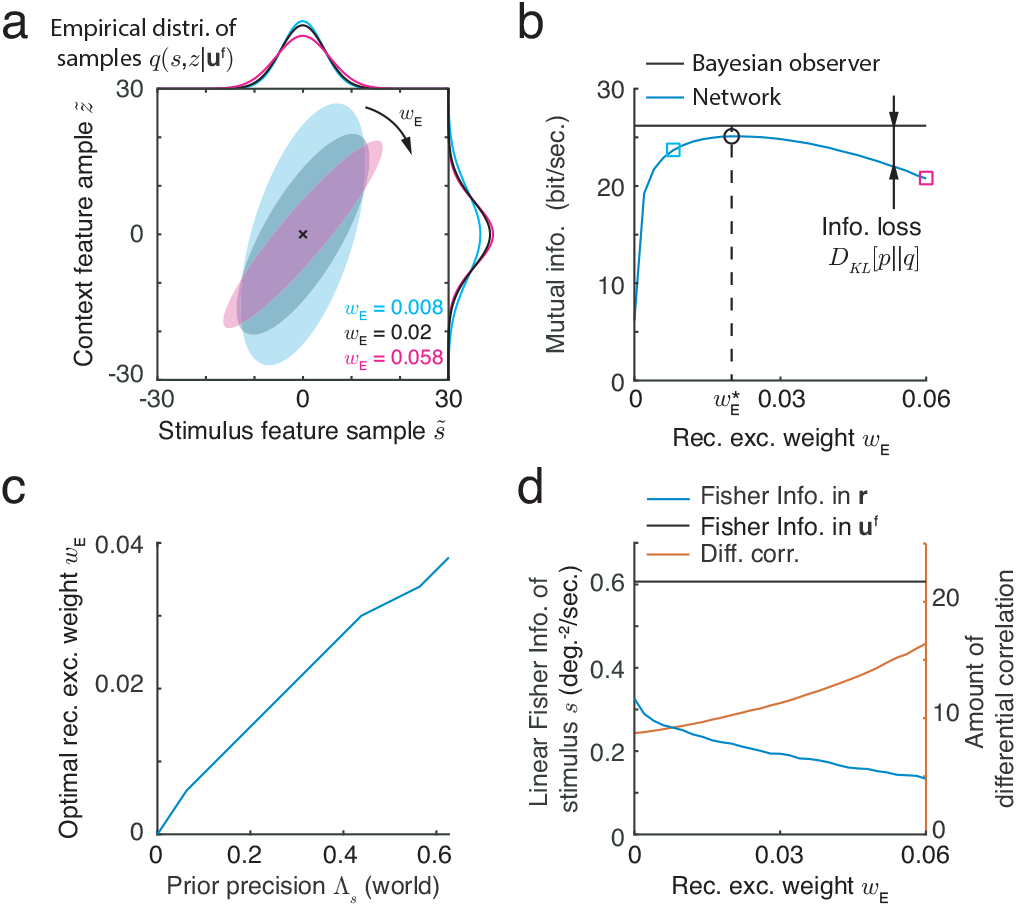
Generating samples from the joint posterior over latent features requires that a network stores an appropriate internal model in its recurrent weights. (a) The joint empirical distribution of samples read out from the population responses for different values of peak recurrent excitatory weight, *w_E_*. The ratio of excitatory and inhibitory weights was fixed. Ellipses capture 3 standard deviation from the mean of the joint distribution of samples. Different colors correspond to the three values of *w_E_*, denoted by different symbols in panel b. (b) The mutual information between latent features, *s* and *z*, and feedforward inputs for ideal Bayesian observer (black) and our network models (blue). The difference between the two lines is the KL divergence between the posterior, *p*(*s, z*|**u**^f^) and the distribution of samples generated by the network, *q*(*s, z*|**u**^f^), which quantifies the loss of mutual information in approximating *p* by *q*. The loss of information is minimized when the weights in the recurrent network, *w_E_*, are set so that *q* best matches *p* (black circle). The three dots correspond to the parameters used in panel (a). (c) The optimal recurrent weight, that maximizes the lower bound on the mutual information in the network’s activity increases with prior precision Λ_*s*_ (Fig. 2c). (d) The linear Fisher information (LFI) of stimulus, *s*, in the population response, **r**, decreases with recurrent weight *w_E_* (blue), which is bounded above by the Fisher information in feedforward inputs **u**^f^ (black). The decrease in LFI is due (in part) to a concomitant increase in differential noise correlations (orange) with *w_E_*.

We quantified the distance between the empirical distribution of samples, *q*(*s, z*|**u**^f^), generated by the network and the true posterior, *p*(*s, z*|**u**^f^) (Eq. 14), using the Kullback-Leibler (KL) divergence. The KL divergence quantifies the loss of mutual information, measured in bits, between latent features, i.e., *s* and *z*, and the feedforward inputs **u**^f^ due to approximating the true posterior distribution, *p*, with the sampling distribution, *q* (Eq. 17) ^36^. KL divergence is always non-negative: A zero KL divergence indicates *q* equals *p*, while a positive KL divergence reflects a deviation of *q* from *p* which causes information loss. Our simulations showed that the loss of mutual information in the network is minimized at a unique value of the recurrent weight, *w_E_*, at which the sampling distribution, *q*, best matches the posterior *p* (Fig. 5b). In contrast, the linear Fisher information of stimulus feature, *s*, in the neuronal responses, **r**, decreases monotonically with *w_E_* (Fig. 1d and 5d, left vertical axis). This is due to the increase in correlated variability with stronger recurrent interactions (Fig. 5d, right vertical axis), resulting in a decrease in discriminability of the external stimulus feature *s* from the neuronal response, **r**.

To confirm that the optimal recurrent weight, 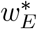, increases with prior precision (Eq. 6), we simulated the network response at different values of Λ_*s*_ in the generative model, and found the weights that minimized the loss of mutual information. The results confirm our theoretical observations (Fig. 5c): When Λ_*s*_ = 0, and the context and stimulus features are uncorrelated, a network with no interactions performs best 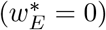. For small Λ *s*(relative to Λ_f_) the optimal weight 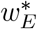 increases roughly linearly with Λ_*s*_, as predicted by our theory.

### Neural signature of sampling in feature subspace: internally generated differential correlations

We next asked what measurable signatures in neural data would characterize sampling based inference of the type we describe. Such sampling would cause instantaneous firing rates, ***λ**_t_*, to fluctuates along the stimulus feature subspace (Fig. 6a), and this would be reflected in covariations of the responses of pairs of neurons within the population. To understand the structure of such covariability, we here describe population response fluctuations along the stimulus feature subspace, since we are mainly interested in how stimulus features are represented in this subspace. For illustration, suppose the mean population response projected onto the feature subspace is centered at *ŝ* = 0° (Fig. 6a). When the population response fluctuates around its mean, the responses of neurons with stimulus preference on the same side of the projected mean position will tend to be positively correlated (the black and blue neurons in Fig. 6a). In contrast, neurons on opposite sides of the mean will tend to be negatively correlated (the black and red neurons in Fig. 6a). A similar argument shows how sampling dynamics structures pairwise correlations in stimulus feature subspace across the entire network (Fig. 6b). We can make these arguments more precise: Our theory suggests that sampling in stimulus feature subspace results in covariability proportional to the product of the derivative of mean population firing rate, i.e., 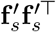 where 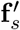 denotes the derivative of mean population rate **f**(*s*) over stimulus feature, *s*. Such correlations have been referred to as differential correlations in previous studies ^2;21^, and they limit the information about *s* that is available to a decoder as measured by linear Fisher Information (see Fig. 1e).

**Figure 6:**
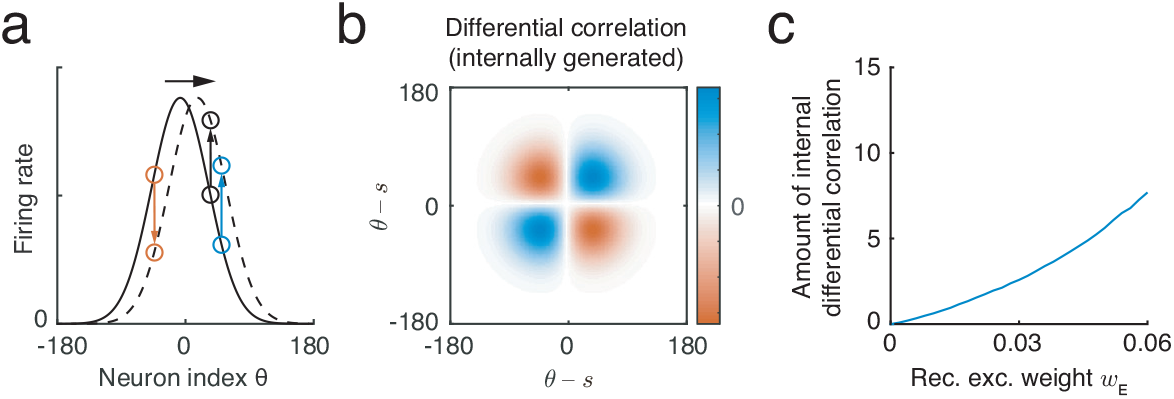
Sampling along the stimulus feature subspace is reflected in internally generated differential correlations. (a) When the network generates samples in the stimulus feature subspace, the population firing rate fluctuates along the feature subspace. Neuron pairs on the same side (the black and blue circles) with respect to the mean of stimulus samples (*s* = 0 in this example), have positively correlated activity, while neuron pairs on opposite sides (the black the red circles) with respect to the mean of stimulus samples are negatively correlated. Fluctuations in the stimulus feature subspace generate correlations proportional to the product of the derivatives of the population firing rates with the stimulus. (b) The pattern of internally generated differential correlation is a result of sampling in the stimulus feature subspace. (c) Internally generated differential correlations increase with recurrent excitatory weight, *w_E_*, which increases with the correlation between stimulus and context in the prior, Λ_*s*_, in a network that samples optimally.

Moreover, in our model, the amplitude of the internally generated differential correlation as a consequence of sampling based inference is determined by the strength of recurrent connections (Fig. 6c). We computed the covariance matrix of the population response, Σ(**r**|**u**^f^), (See Eq. 28), and found that the amplitude of internally generated differential correlations is proportional to the recurrent weight 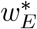,

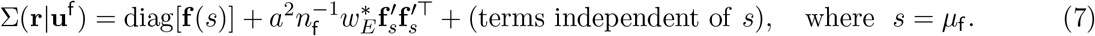

The second term on the right hand side in above equation represents internally generated differential correlations with amplitude 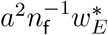, where *a* is the tuning width, *n*_f_ is the total number of feedforward input spikes received by all E neurons (Table 1). diag[**f**(*s*)] is a diagonal matrix capturing the individual Poisson spiking variability, with **f**(*s*) the tuning of E neurons over *s*. Since the optimal recurrent weight, 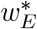, is determined by the correlation between stimulus and context in the prior (Fig. 5c, Λ_*s*_ in Eq. 6), the amplitude of differential correlations in a network implementing optimal sampling is determined by the correlations between stimulus and context features, Λ_*s*_ (Fig. 5d and Eq. 29). We confirmed this conclusion in network simulations which showed that differential correlations increased with excitatory recurrent weight, *w_E_* (Fig. 6c), and hence with Λ_*s*_ (Fig. 5c).

**Table 1:** Notations in the model

| Symbol | Description |
| --- | --- |
| $z$ | Context feature (scalar variable) |
| $s$ | Stimulus feature (scalar variable) |
| $\Lambda_s$ | The prior precision |
| $\mathbf{r}_t$ | Neuronal spikes at time $t$ (vector) |
| $\boldsymbol{\lambda}_t$ | Firing rate at time $t$ (vector) |
| $\mathbf{u}^f$ | Feedforward spiking inputs (vector) |
| $\mathbf{u}_t^r$ | Recurrent input at time $t$ (vector) |
| $U^f$ | Peak feedforward input rate |
| $U^r$ | Peak value of recurrent input |
| $n_f$ | The total spike count of $\mathbf{u}^f$ to all neurons (scalar variable) |
| $n_r$ | The sum of recurrent input (scalar variable) |
| $w_E$ | The peak recurrent weight between excitatory neurons |
| $\theta_j$ | The preferred stimulus feature of $j$-th excitatory neuron |
| $a$ | Tuning width |

To understand why internally generated differential correlations are shaped by the structure of the generative model, we consider the case of uncorrelated context and stimulus (Λ_*s*_ = 0), as we did when examining linear Fisher information (Fig. 1a). In this case, an optimal network should have no recurrent connectivity, and instantaneous firing rates, ***λ****_t_*, will be fixed in response to a fixed feedforward input. We find that in this case the variability in the population response, **r**, is only due to independent Poisson variability in spike generation (the 1st term on the right hand side of Eq. 7). Increasing the correlations between stimulus and context (Fig. 6c) results in larger recurrent weights, 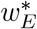 (Fig. 6c), which increases recurrent variability (Fig. 1d, red), and hence increases internally generated differential correlations, thus reducing linear Fisher Information.

### Distinguishing external and internal differential correlations

In many neurophysiology experiments a stimulus feature, *s*, is fixed, while the feedforward input, **u**^f^, fluctuates (Fig. 4a). Hence in experimentally recorded neuronal responses, fluctuations in the stimulus feature subspace reflect a combination of sensory transmission noise, and any internally generated variability. Differential noise correlations in the evoked population activity, **r**, in response to a fixed stimulus, *s*, can be decomposed into external correlations inherited from fluctuations in **u**^f^, and the correlations internally generated from recurrent network interactions. In this case the total magnitude of differential correlations in the evoked response is 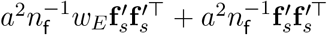 (Eq. 30), where the new term, 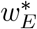, reflects differential correlations inherited from the input (compare with Eq. 7). Although the two sources of differential correlations are intertwined, we found that they impact information processing differently. This difference offers a potential way to distinguish between them, and determine whether the network produces samples in the joint stimulus and context feature subspace.

Externally imposed differential correlations decrease with the strength (i.e. contrast) of the feedforward input (the peak firing rate of feedforward input, Eq. 10) (Fig. 7a, orange curve). Consequently, the LFI of *s* in the observations of either the feedforward input, **u**^f^, or the population activity, *r*, increase with the strength of the feedforward input (Fig. 7a, black and blue curves, respectively). Similarly, the mutual information (the information between feedforward inputs **u**^f^ and the latent features, i.e., *s* and *z*, sampled by the network, Eq. 17 and Fig. 5b) also increases with feedforward input intensity (Fig. 7b). We therefore have a monotonic, decreasing relationship between externally generated differential correlations and mutual information (Fig. 7c). This is expected since such inherited correlations always impair information processing, as observed previously ^2;21^. The magnitude of internally generated differential correlations is proportional to the recurrent excitatory weight, *w_E_* (Eq. 7). The non-monotonic relation between *w_E_* and the mutual information between **u**^f^ and *s* and *z* (Fig. 5b) creates a similar non-monotonic relation between internally generated differential correlations and mutual information (Fig. 7d). Therefore, the impact of external and internal differential correlations on stimulus coding can be distinguished by their respective monotonic and non-monotonic relation with the mutual information in the network (see Discussion).

**Figure 7:**
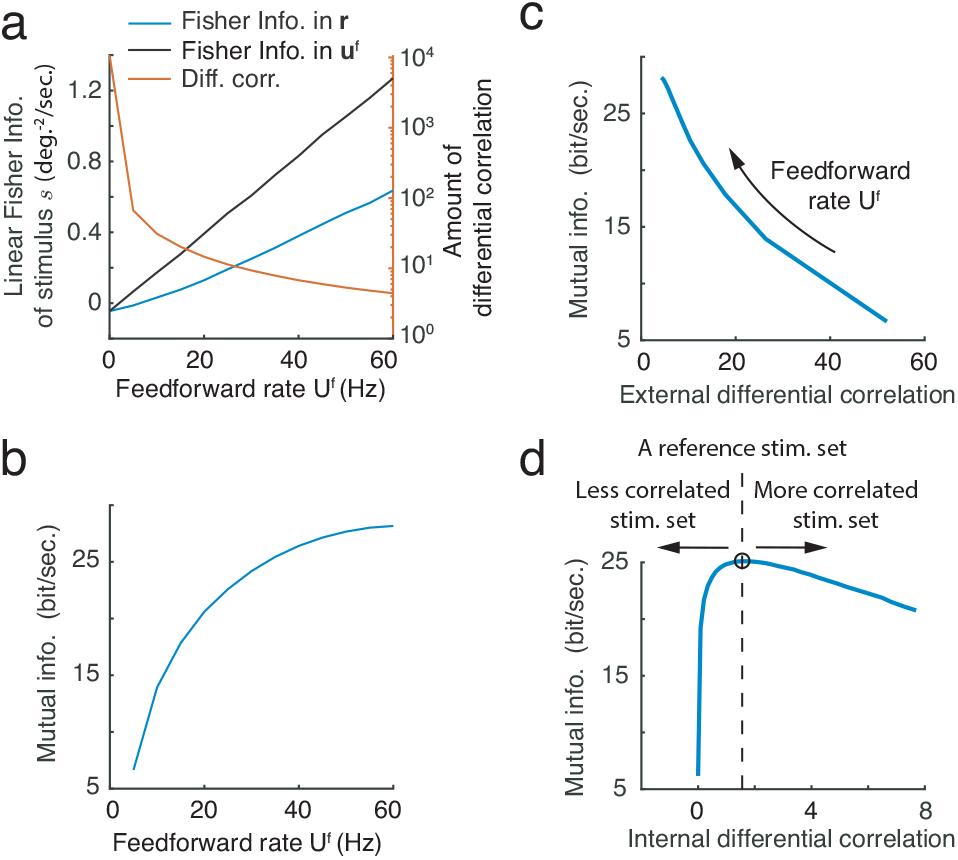
(a-b) Externally generated differential correlations, inherited from feedforward inputs, impair information representation. (a) The feedforward rate determines the input gain, and the reliability of sensory transmission and the amount of external differential correlation inherited from feedforward input. (b) Mutual information (the information between the feedforward inputs and the latent features sampled by the network with optimized recurrent weight 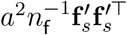) monotonically increases with feedforward rate. (c-d) External and internal differential correlations have a distinct impact on information representation in the recurrent circuit model. While we find a monotonic relation between mutual information in a recurrent network and external differential correlations (c), there is a non-monotonic relation between mutual information and internal differential correlations (d). The direction of the arrow denotes the change of the parameters that lead to the change in differential correlations.

## Discussion

In this study we propose a new role for recurrent connections in neural circuits that represent complex sensory scenes. We demonstrated their function in a model cortical hypercolumn that implements a sampling-based approximation of the joint posterior over a local stimulus and a global context feature. There is strong evidence that Bayesian inference underlies perceptual processing ^12;13;15;32^, such as multisensory cue integration^11^ and sensorimotor learning ^14^. In our circuit model, the recurrent connections store an associative prior between the stimulus and context features, and thus reflect the statistical structure of the world learned (or assumed) by the animal. Internally generated Poisson variability, reflecting the known trial-to-trial and dynamic variability in neural dynamics, provides the stochasticity needed for implementing the sampling algorithm. Finally, the stimulus and context features are embedded in a subspace in the neural population responses from which the posterior of features can be linearly read out.

A neural signature of network sampling in stimulus feature subspace is the covariability of population responses along the stimulus feature subspace, which is a reflection of internally generated differential noise correlations. Such correlations are due to recurrent interactions in the network, and not inherited from sensory transmission noise in feedforward inputs^21^. We have shown that the amplitude of internally generated differential correlations in networks that are tuned to accurately sample from the posterior distribution are determined by the associative prior between stimulus and context features. This observation could be used to verify whether cortical circuits produce samples from a posterior distribution over the stimulus features, as we discuss below. While externally and internally generated differential correlations are necessarily intertwined in any recorded neuronal responses, we found that they impact information representation in the circuit differently. The mutual information between feedforward inputs and latent features sampled by the network is maximized at a unique amplitude of internal differential correlations (Fig. 5 and 6), while external differential correlations always decreased mutual information and hence impaired information processing (Fig. 7c).

The proposed circuit sampling mechanism thus produces experimentally testable predictions. First, our model requires that the feedforward inputs and the recurrent inputs have the same tuning profile (Eq. 1 and 3), in order to compute the conditional distribution of latent stimulus feature via summing feedforward and recurrent inputs (Eq. 4). The assumption that feedforward and recurrent inputs have the same tuning profile is supported by a recent circuitry experiments in layers 4 and 2/3 in mouse V1 ^9^. Second, our model predicts that the magnitude of internal differential correlations is determined by the learned correlation between stimulus and context (Fig. 6b-c). This could be tested with a perceptual learning task in which a subject is repeatedly presented with the same set of stimuli ^43^. We predict that increasing the correlation between global context and the local stimulus in the stimulus set, would lead to an increase in differential correlations during and after learning (See Fig. 5c-d). Indirect evidence from previous studies supports the possibility that neuronal correlations and spatiotemporal firing patterns are shaped by external stimuli: It has been shown that neurons become less correlated with perceptual learning with a simple stimulus set ^44^, and spontaneous population response patterns, including correlation structure, become more similar to evoked response patterns during development^45^. Lastly, to verify the predicted impact of internal differential correlations on information representation, we could test whether there exists a non-monotonic relation between the magnitude of internally generated differential correlations and the mutual information (the information between feedforward inputs and latent features sampled by the network) estimated from population responses (Fig. 7d). For example, suppose an animal is fully adapted to a *reference* stimulus set with a prescribed correlation between the context and local stimulus. Training the animal with a more or less correlated stimulus set (compared to the reference set) would then increase or decrease the amplitude of internal differential correlations respectively. Regardless of whether the differential correlations increase or decrease, our network model predicts that mutual information (with the true generative model, *p*(Θ) in Eq. 17, defined as the reference stimulus set) in the population responses will decrease (Fig. 7d).

The recurrent circuit model we presented includes novel mechanisms for inference and representation, which combine features from previously proposed neural coding frameworks, including parametric representations characteristic of probabilistic population codes (PPCs)^16;34^, and features of sampling-based codes (SBCs) ^17;19;37;38;40^. In our network, the conditional distribution of the stimulus feature is represented by instantaneous firing rates using a parametric code similar to a PPC (Eqs. 3 and 4), i.e., the conditional distribution belongs to a linear exponential family of distributions with sufficient statistics that are linear projections of instantaneous firing rates. Further, our network produces samples in the stimulus feature subspace with variability generated internally, similar to previously proposed SBCs (Fig. 3a).

Despite these similarities, there are some fundamental differences between our network and PPCs and conventional SBCs. PPCs are generally implemented in deterministic networks (no internal variability) that sum Poissonian inputs, and variability of neuronal responses is inherited from feedforward inputs. In contrast, the network model we proposed is doubly stochastic and includes both internal Poisson variability that drives sampling in the joint stimulus and context feature subspace, as well as variability in feedforward inputs to the network which conveys the likelihood function ^34^ (Fig. 3a). Second, in PPCs the posterior of stimulus feature is represented parametrically in neuronal responses, while in our proposed network the joint posterior over stimulus and context features is approximated by sequential samples. Most conventional SBCs rely on the assumption that the profile of averaged population responses is directly proportional to the posterior over a latent stimulus feature ^17;19;20;37;38;40^, i.e., (**r**) ∝ *p*(*s*|**u**^f^), while in our network the posterior is approximated by samples in the stimulus feature subspace in the space of neuronal population responses. The samples of stimulus and context features can be read out linearly from the instantaneous spiking activity, **r**_*t*_, and recurrent input 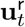 respectively (Fig. 4). Hence, in our proposed network the sampling space has dimension two, while in conventional SBCs the dimension of sampling space is the number of neurons in the network. A benefit of sampling in the feature subspace is that sampling is much more efficient in spaces of lower dimension.

On the network level, the recurrent connections are translation-invariant in the stimulus feature subspace, an assumption widely used in continuous attractor network models (CAN) ^26;46–48^. Translation-invariant connections are a mathematical simplification, but are not required for a circuit to implement sampling. CANs have been shown to achieve maximal likelihood estimation via template matching^16^. Here we have shown that a network with CAN-like structure and internally spiking variability is able to perform sampling in stimulus feature subspace to approximate the posterior over stimulus features, rather than providing a point estimate as in most deterministic CANs. As Poissonion spiking is ubiquitous in cortex ^4^, real cortical circuits may use it to sample stimulus features.

To keep the exposition transparent we have presented a model of minimal complexity. The proposed network mechanism of sampling-based inference can be generalized to more complex generative models, and hence the assumption of Gaussianity (Eqs. 8 and 9) and analytical expression in Eq. (14) are not essential to the proposed mechanism. For example, a similar network can generate samples from other types of distributions belonging to the linear exponential family^34;36^. The type of distribution from which the network samples is determined by the basis function (logarithm of tuning) **h**(*s*) (Eq. 4). These basis functions depend on the tuning of feedforward inputs, along with the recurrent connection profiles, and do not have to be Gaussian. The network model could also infer the latent features with a non-uniform prior, when the preferred stimuli of neurons in the population are not distributed uniformly in the stimulus feature subspace. In addition, the network could also produce samples from a high-dimensional stimulus feature distribution, (*s*_1_, *s*_2_,…, *s_m_*). In this case, features could be inferred jointly by coupled networks consisting of neurons with Poissonian spiking variability, with each subnetwork inferring the marginal posterior of one feature. The recurrent connections in the coupled networks then need to be set to make the mean recurrent inputs received by each network have the same profile as the feedforward inputs received by that network. Our recent preliminary study demonstrates this possibility ^49^.

In conclusion, we have shown that recurrent connections in a cortical circuit can store information about the learned structure of the world, and that neural noise can drive sampling from the posterior over directly and indirectly observed features. Internally generated differential correlation are a signature of the sampling of stimulus features, and the size of these correlations is determined by the representation of the structure of the world stored in recurrent circuits.

## Methods

### The probabilistic generative model

We assume that the statistics of stimuli in the world, and the feedforward inputs received by neurons in the recurrent network, can be described by a hierarchical generative model. For illustration, we use a hypercolumn of V1 neurons as a concrete example to explain the underlying idea (Fig. 2a), although our generative model can be generalized to explain information processing in other brain areas. We denote by *z* the one dimensional context feature (e.g., orientation) of an extended object in the visual field, which represents the contextual information and may cover the receptive fields of multiple V1 hypercolumns. Although the orientation distribution of natural images is bimodal, we assume that *z* follows a uniform distribution for simplicity (Fig. 2c, marginal plots),

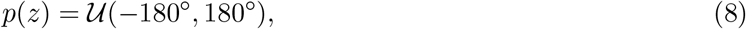

where 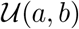 denotes a uniform distribution over [*a, b*]. The assumption that the prior distribution over the context variable is uniform simplifies our model significantly, as it implies the spatial homogeneity of the network model as given by Eqs. (10–11). However, it doesn’t substantially change our main results. We denote by *s* the one dimensional feature (e.g., orientation) of the local stimulus falling into the central receptive field (cRF) of a single V1 hypercolumn. Due to the differences between the local (stimulus) and global (context) aspects of the sensory scene, the stimulus feature *s* is not identical to the contextual feature *z*, but we assume that the two are related as,

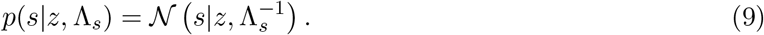

Here 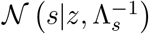 denotes a normal distribution over *s* with mean *z* and precision (inverse of variance) Λ_*s*_.

Given the stimulus feature *s* located in the center receptive field of the V1 hypercolumn, we model the feedforward inputs **u**^f^ to the *N_E_* excitatory neurons in V1 as conditionally independent Poissonan variables, with Gaussian intensity tuning over the input,

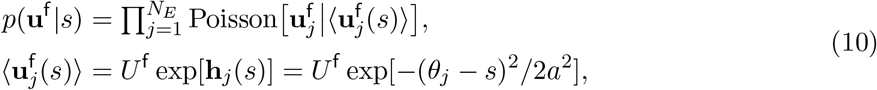

where 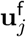 denotes the feedforward input received by *j*^th^ excitatory neuron, 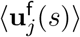 is the tuning of the feedforward input, with *θ_j_* denoting the preferred stimulus feature of 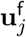, and *U* ^f^ and *a* characterize the peak firing rate and width of the tuning function, respectively. This model of the input has been widely used in previous neural coding studies, such as those of probabilistic population codes (PPC) ^16;34;35^.

Under the assumption that the preferred stimulus feature values, 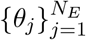, of all feedforward inputs are uniformly distributed in the space of the stimulus feature *s*, we can use Eq. (10) to show that the likelihood of *s* given a single observation of the input, **u**^f^, belongs to the exponential family of distributions with linear sufficient statistics ^16;34;35^, and has the form

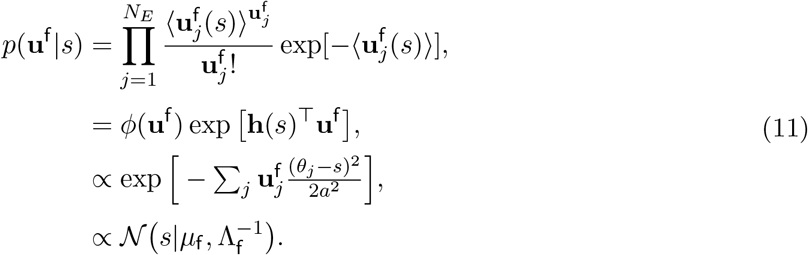

The second line Eq. (11) denotes the standard form for a distribution belonging to the linear exponential family, where *φ*(**u**^f^) is a function which only depends on the input, **u**^f^. The *basis function*, **h**(*s*) ^16^, determines the profile of the input tuning (Eq. 10), as well as the type of likelihood distribution (Eq. 11). Since the input tuning has a Gaussian profile, the likelihood of *s* given **u**^f^ can be normalized to obtain a Gaussian distribution (Eq. 11), with mean *μ*_f_ and precision Λ_f_ which are both linear functions of the inputs, 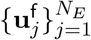,

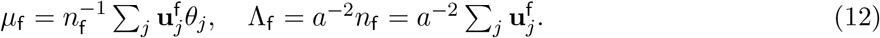

The mean, *μ*_f_, represents the position of **u**^f^ in stimulus feature subspace, and the precision, Λ_f_, is proportional to the sum of total feedforward spike counts, *n*_f_ (Fig. 2d-e). In sum, the whole generative model is determined by the joint distribution,

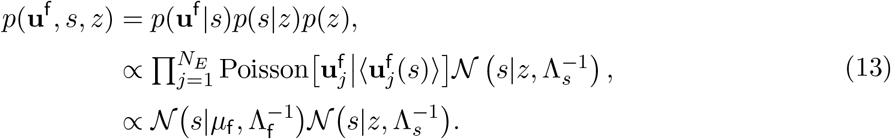

### Approximate Bayesian inference via Gibbs sampling

We assume that the neural circuit needs to compute information about both the local feature, *s*, and context feature, *z*, from the feedforward input **u**^f^. The circuit approximates the joint posterior distribution over the features, rather than making a point estimate ^12;32^. The simplicity of the generative model (Eq. 13), allows us to obtain an analytical expression for the posterior distribution,

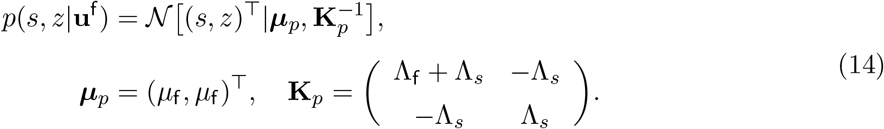

Our goal is to develop a biologically plausible sampling algorithm to approximate this posterior distribution, and use Eq. (14) to verify the output of the algorithm.

We consider using Gibbs sampling ^36^ (a type of Monte Carlo method) to approximate the posterior. To describe the iterative Gibbs algorithm, we assume that a sample, 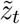, of the context feature, *z*, is provided at time *t*. We can use this sample, 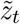, with the feedforward input, **u**^f^, to update the conditional distribution of stimulus feature *s* (step 1 in Fig. 2f),

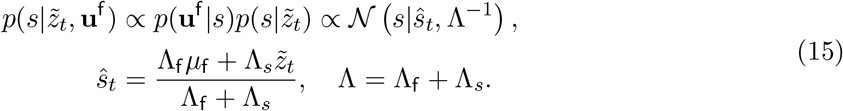

Here *ŝ*_*t*_ and Λ denote the mean and precision (inverse of variance) of the conditional distribution 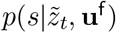. The next step in the algorithm is to draw a sample, 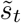, from the conditional distribution 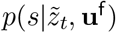 (step 2 in Fig. 2f),

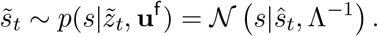

Next, the conditional distribution over the context feature, *z*, is updated given this new sample, 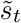, and a new sample, 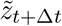, is drawn (step 3 in Fig. 2f),

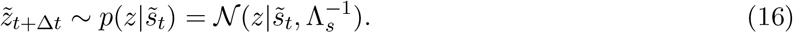

These three steps in the Gibbs sampling algorithm (Eqs. 15–16) are performed iteratively until the distribution of samples, 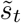 and 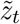, approximate the true posterior distribution with sufficient accuracy (Fig. 2g; compare the red dots with the blue contour map).

### Inference from an information-theoretic point of view

The goal of the sampling algorithm is to approximate the posterior distribution over the features, Θ = {*s, z*}, given a feedforward input, **u**^f^. When the sampling algorithm uses a model which does not match the structure of the generative model for the stimuli, the samples will converge to distribution *q*(Θ|**u**^f^) which differs from the true posterior, *p*(Θ|**u**^f^) (Eq. 14). In this case the mutual information between the asymptotic samples of latent features, Θ, and feedforward input will be smaller than in the case when samples come from the true posterior, *p*(Θ|**u**^f^),

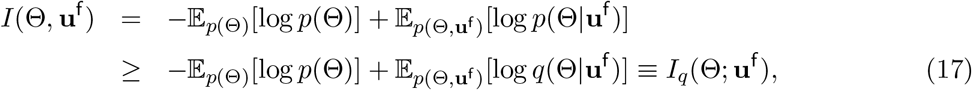

It is straightforward to show that the difference between *I*(Θ, **u**^f^) and *I_q_*(Θ, **u**^f^) is the Kullback-Leibler (KL) divergence between *p* and *q*, i.e., 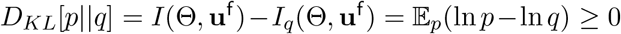. Equality in Eq. (17) holds only if the distribution *q* matches the true posterior *p* exactly.

The mutual information *I_q_*(Θ;**u**^f^) can be computed analytically when the approximating distribution 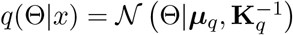 is a bivariate normal (substituting Eqs. 13 and 14 into Eq. 17),

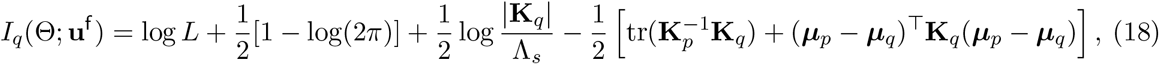

where *L* = 360° is the length of the stimulus feature subspace. Here ***μ****_p_* and **K**_*p*_ are the mean and the precision matrix of the posterior distribution (Eq. 14). When *q* matches the posterior distribution, *p*, we have, 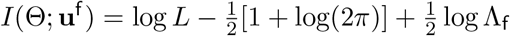. As expected mutual information increases with the reliability of sensory transmission, Λ_f_ (Fig. 7b).

### A stochastic recurrent network sampling latent features

We consider a generic recurrent network consisting of *N_E_* excitatory Poisson neurons and implement sampling from the posterior over the features using network dynamics. For simplicity, we describe neuronal activity using a Hawkes process (a type of inhomogeneous Poisson process^50^). The instantaneous firing rates of the neurons in the network at time *t*, ***λ***_*t*_, obey the following recurrent equations:

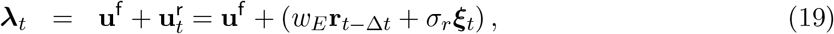

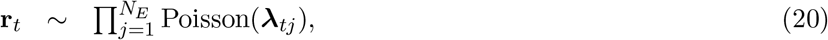

where **u**^f^ is the feedforward Poisson spike input (Eq. 10), 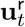 is the continuous valued recurrent input at time *t*. At each time, the neurons independently generate Poisson distributed spike counts **r**_*t*_ (over the interval [*t* − Δ*t, t*]) from the instantaneous rate ***λ***_*t*_. The parameters *w_E_* and *σ_r_* denote the excitatory recurrent weight and recurrent variability, respectively. Below we present the conditions for the network to implement Gibbs sampling.

### Representation in neuronal activity

We require the network to generate and maintain a representation, in which the recurrent input, **u**^r^, represents the context feature, *z*, and the activity, **r**represents the stimulus feature, *s*. Specifically, we assume that with fixed spiking activity **r**_*t*−Δ*t*_, which represents stimulus feature sample 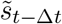, the recurrent inputs, 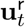, have independent Poisson-like variability (the variance equals the mean), and its mean, 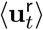, has a Gaussian profile over 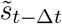,

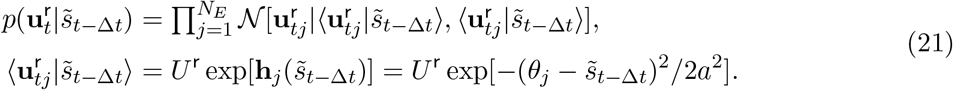

In Eq. (21) we also assume that the recurrent input, **u**^r^, has the same tuning profile as the feedforward input (Eq. 10) (by sharing the same basis function **h**(*s*)), and hence the recurrent and feedforward inputs received by the same *j*^th^ neuron have the same preferred stimulus *θ_j_*. The assumption of independence of the components of **u**^r^ given a stimulus feature sample, 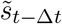, does not mean that the components of **u**^r^ conditioned on an external stimulus feature *s* are independent. Due to the nature of the sampling dynamics the recurrent input **u**^r^ fluctuates in the stimulus feature subspace, correlating activity through an external stimulus feature (Eq. 7).

Since we consider a network that transforms its inputs linearly (Eq. 19), if we only consider the linear Fisher information of *s* contained in **u**^r^, then the likelihood of *s* given a recurrent input **u**^r^ can be normalized to obtain a Gaussian distribution as follows,

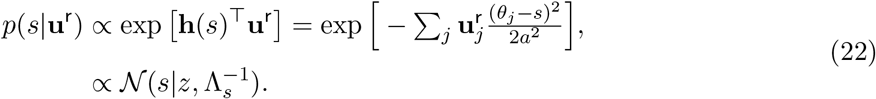

Comparing Eqs. (22) and (9) we see that the recurrent input **u**^r^ can represent a conditional distribution over the stimulus feature *s*, and that the mean and precision of the prior can be linearly read out from **u**^r^,

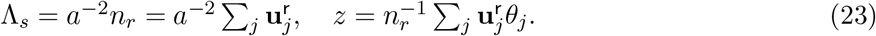

Because the recurrent and feedforward inputs represent the distributions in the same way by design (see Eqs. 11 and 22), the sum of feedforward and recurrent inputs gives the instantaneous firing rate ****λ***_t_* (Eq. 19),

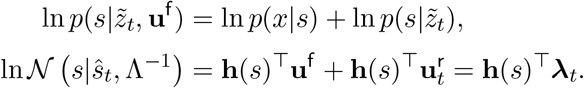

Therefore the instantaneous firing rate, ***λ****_t_*, represents the conditional distribution 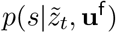, with the precision and the mean of the conditional distribution determined by linear functions of the rates, Λ = *a*^−2^Σ_*j*_**λ**_*tj*_ and 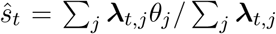. Table 2 lists how the population inputs, and instantaneous firing rate in the recurrent network represent the parameters of the distributions. The average (tuning) of ***λ*** can be computed as,

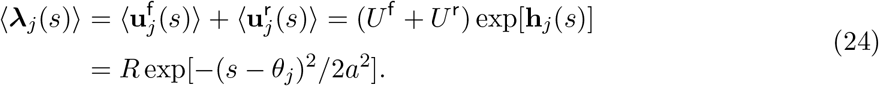

**Table 2:**
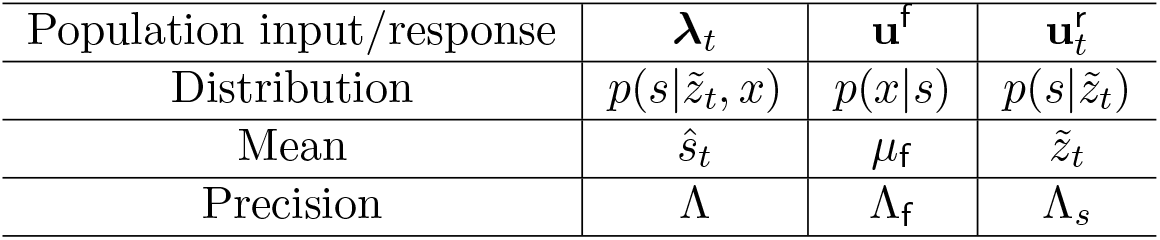
Representation in neural population inputs and responses.

### Poisson spike generation samples stimulus feature subspace

We demonstrate next that the internally generated Poisson spike variability generates random samples from the conditional distribution 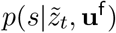 of the stimulus feature, *s*, as long as the instantaneous firing rate, ***λ****_t_*, represents 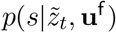 as in Eq. (4). Specifically, we show that the empirical distribution of the stimulus sample, 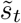, read out from **r**_*t*_ is consistent with 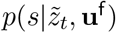 represented in ***λ***_*t*_. To simplify the derivation, suppose the time window or neural population are sufficiently large so that the instantaneous firing rate, ***λ_t_***, is close to its trial average, i.e., a smooth Gaussian profile (Eq. 24), so that 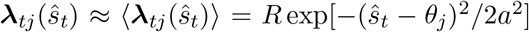. When generating Poisson spikes **r**_*t*_ from 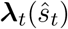 (Eq. 20), the likelihood of an emitted spike **r**_*t*_ given *ŝ*_*t*_ is proportional to a Gaussian distribution,

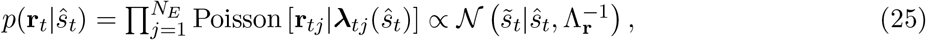

with precision Λ_*r*_ = *a*^−2^*n_r_* = *a*^−2^Σ_*j*_**r**_*tj*_ and 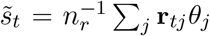. Comparing the empirical distribution of stimulus feature sample 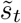 collected over trials (Eq. 25), and the conditional distribution 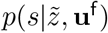 (Eq. 4), we see they are both Gaussian distributions with the same mean *ŝ*_*t*_ but different precision (Λ_*r*_ versus Λ). However, the precision of the empirical distribution, Λ_*r*_, equals the precision of conditional distribution, Λ, in the limit of a large neural population, or a long time window (see Supplementary Information for a rigorous derivation),

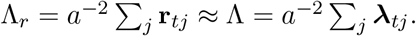

This shows the Poisson spike generation leads the neural dynamics to asymptotically sample stimulus features from the true conditional in the limit of a large neural population, or over long time windows. We confirm this conclusion in simulations (Fig. 3h).

### Recurrent connections store the association between stimulus and context

The representation of context feature samples in recurrent input, **u**^r^, (Eqs. 21 and 22) puts constraints on the recurrent weight, *w_E_*, and recurrent variability, *σ_r_*. Because the stimulus tuning of the recurrent input, **u**^r^, and responses, **r**, have the same width *a* (Eqs. 21 and 24), every neuron in the population can only be connected to itself in the simple network consisting of only excitatory neurons, since otherwise the tuning of **u**^r^ would be wider than that of **r**. Combining Eqs. (21 and and using Table 2, we can show that every neuron is self-connected with the same weight *w_E_*, with the optimal weight satisfying the requirement of recurrent inputs (Eq. 21),

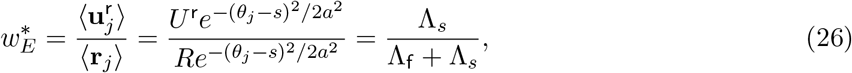

which yields Eq. (6) in the main text. The last equality in Eq. (26) follows from the observation that the sum of the vector components **r**and **u**^r^ represent the precision of the conditional distribution, *p*(*s*|*z*, **u**^f^) and the prior *p*(*s*|*z*), respectively (Table 2). The self-connections are a result of the simplifying assumption that the network consists solely of excitatory neurons (Eq. 19). This assumption simplified the analysis, and is relaxed below (see Eq. 33 and Fig. 1).

Another assumption on the recurrent connections is that the components of **u**^r^ exhibit independent, Poisson-like variability (the variance is the same as mean), given the network’s activity at the previous time step, **r**_*t*−Δ*t*_. This constrains the recurrent variability *σ_r_* (Eq. 19). Hence,

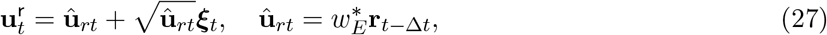

where ***ξ***_*t*_ is a *N_E_* dimensional Gaussian white noise with 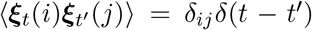, where *δ_ij_* and *δ*(*t* − *t*′) are Kronecker and Dirac delta functions respectively. The Poisson-like variability in recurrent interactions allow for the sampling of the context feature (Eq. 2c), via the same mechanism that allowed for sampling of the stimulus feature using Poisson spike generation (Eq. 5).

### Analysis of the conditional distribution of the neuronal response

We compute the distribution of neuronal responses, **r**, given an external stimulus feature, *s*, which can be compared with experimental observations ^22^. For a fixed external stimulus, *s*, the neuronal response, **r**, fluctuates over time/trial due to both sensory transmission noise described by *p*(**u**^f^ |*s*) (Eq. 13 and Fig. 2b), as well as the internally generated variability described by *p*(**r**|**u**^f^) (Fig. 3a). Together, the distribution of **r**in response to an external stimulus *s* therefore has the form

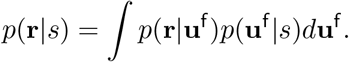

For simplicity, we only compute the covariability of *p*(**r**|**u**^f^) along the stimulus feature space, since we are mainly interested in how stimuli are represented in this subspace. By approximating the Poissonian spiking variability *p*(**r**|****λ****) by a multivariate normal distribution (Eq. 20), and considering the limit of weak fluctuations in ****λ**** along the stimulus feature subspace, *p*(**r**|**u**^f^) can be computed approximately as (see math details in Supplementary Information),

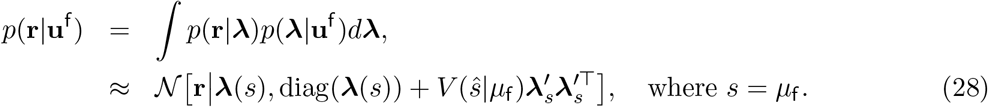

The covariance structure of the neuronal response includes two terms: diag(***λ***(*s*)), a diagonal matrix whose entries equal that of the vector ***λ*** denoting the (independent) Poisson spiking variability (Eq. 13), and 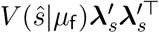, a term that captures the covariability due to firing rate fluctuations along the stimulus feature subspace, where 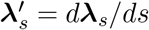 is the derivative of ***λ****_s_* over the stimulus feature *s*. The covariance 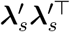 is often termed differential correlations ^2;21^. With the Gaussian tuning of ***λ****_s_* (Eq. 24), 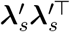 exhibits anti-symmetric structure (Fig. 6b), as we noted previously ^21;26;51–53^.

In Eq. (28), *V* (*ŝ*|*μ*_f_) characterizes the amplitude of internal differential correlations, and *ŝ* and *μ*_f_ denote the position of ****λ**** and **u**^f^ on the stimulus feature subspace respectively (Table 2). The dynamics of Gibbs sampling (Eq. S5 in Supplementary Information) and the network structure (Eq. 6) imply that

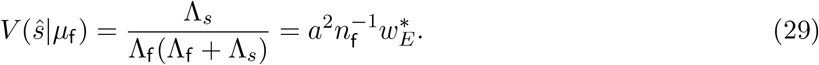

Note that *V* (*ŝ*|*μ*_f_) is constrained by network connections, in that it is internally generated by the network. Consequently, any internally generated differential correlations can be regarded as a neural signature of the network performing sampling in stimulus feature subspace.

An expression for *p*(**r**|*s*) can be derived similarly, and includes an additional term contributing to differential correlations compared with *p*(**r**|**u**^f^) (Eq. 28) due to fluctuation in the feedforward inputs,

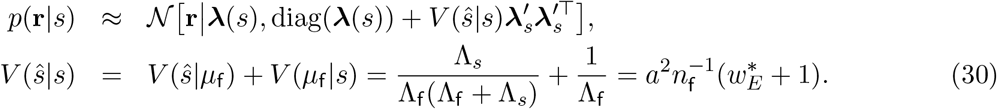

Here the variance *V* (*ŝ*|*s*) in the stimulus feature subspace is a mixture of internal variability, *V* (*ŝ*|*μ*_f_), and sensory noise, *V* (*μ*_f_ |*s*) (Eq. 13).

### A spiking network model with excitatory and inhibitory Poisson neurons

To test the validity of the proposed mechanism we developed a recurrent network model consistent with many previous mechanistic models of cortical dynamics ^25;26^. The network consists of *N_E_* excitatory (*E*) and *N_I_* inhibitory (*I*) spiking neurons, with the activity of each neuron modeled as a Hawkes process ^50^. At time *t*, we represent the response of neuron *j* in population *a* = {*E, I*}, 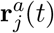, as a spike count drawn from a Poisson distribution with instantaneous firing rate, 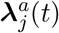,

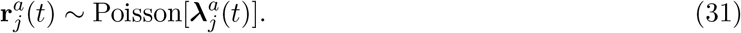

The firing rate 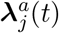 is the sum of feedforward input 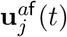 and recurrent input 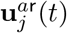, so that 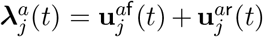. The feedforward inputs are filtered spikes from upstream neurons, 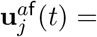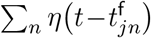, where 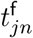 is the time of the *n*-th spike received by neuron *j* of type *a* from feedforward inputs. Here *η*(*t*) is the synaptic input profile which is modeled as *η*(*t*) = exp(−*t/τ_d_*)*/τ_d_*, (*t* >0). Throughout, we set the synaptic time constant *τ_d_* = 2ms. The recurrent input received by neuron *j* in population *a* is defined by

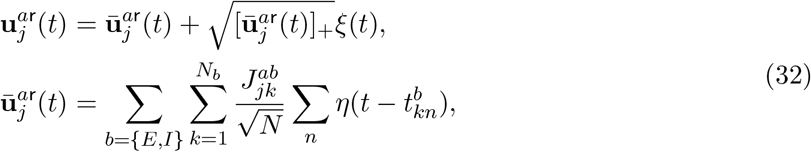

where 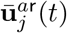 is the mean recurrent input at time *t* given the neuronal activities of the presynaptic neurons. The recurrent input in the network is corrupted by noise whose variance equals the mean of the recurrent input. In a physiological network, recurrent noise may be generated by the chaotic state in network dynamics ^54^ or synaptic noise ^41;42^. In Eq. (32) the function [·]_+_ rectifies the negative input, and *ξ*(*t*) is a random varialbe following a standard Gaussian distribution. The coefficient 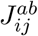 is the synaptic weight from neuron *j* in population *b* to neuron *i* in population *a*. The time 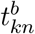 is the time of the *n*^th^ spike fired by neuron *k* in population *b*. The parameter *N* = *N_E_* +*N_I_* is the total number of neurons in the network. The scaling of the synaptic weights by 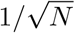 is standard in networks where excitation is balanced by recurrent inhibition ^54^. Finally, the synaptic input profile of the recurrent input, *η*(*t*), is the same as the one we chose for the feedforward input for convenience.

To model the coding of a circular stimulus feature such as orientation, the excitatory neurons are arranged on a ring ^26;51^. The preferred stimuli, *θ_j_*, of the excitatory neurons are equally spaced on the interval [−180°, 180°], consistent with the range of latent features (Eq. 8). Inhibitory neurons are not tuned to stimulus orientation, and their role is to stabilize network responses. The recurrent connections between *E* neurons are modeled using a Gaussian function decaying with the distance between the stimuli preferred by the two cells,

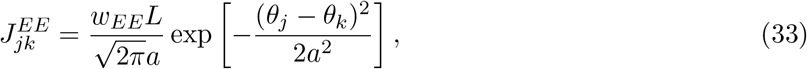

We imposed periodic boundaries on the Gaussian function to avoid boundary effect in simulations. Although in the generative model we assumed non-periodic feature variables (Eq. 1), as long as the variance of the associated distributions are smaller than the width of the feature space, the network model with periodic boundaries on the recurrent connection (Eq. 33) provides a good approximation of the non-periodic Gaussian posterior (Eq. 14). The weight *w_EE_* denotes the average connection strength of all *E* to *E* connections. The parameter *a* = 40° defines the footprint of connectivity in feature space (i.e the ring), and *L* = 360° is the length of the ring manifold (Eq. 8); Multiplication by *L* in Eq. (33) sets the sum of all *E* to *E* connection strengths equal to *N_E_w_EE_*. Moreover, the excitatory and inhibitory neurons are all-to-all connected with each other (similar for *I* to *I* connections). For simplicity, we consider the *E* to *I*, *I* to *I* and *I* to *E* connections all to be unstructured (in feature space) and assume that connections of the same type have equal weight, i.e.,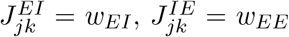 and 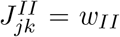. To simplify the network further, we consider the connections from the same population of neurons to have the same average weight, i.e., *w_EE_* = *w_IE_* ≡ *w_E_* and *w_II_* = *w_EI_* ≡ *w_I_*.

The feedforward inputs applied to E neurons consist of independent Poisson spike counts as described by Eq. (10), with rate 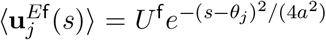. The inhibitory neurons also receive feedforward indpendent Poissonian inputs. The firing rate of the input received by every *I* neuorn is proportional to the overall feedforward rate of input to *E* neurons, in order to keep the excitatory and inhibitory balance of neuronal activities in the network,

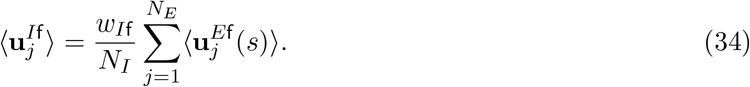

In the simulations shown in Figs. 4–7, we started with a network of *N_E_* = 180 excitatory and *N_I_* = 45 inhibitory neurons, and increased the number of neurons by a fixed factor in Fig. 1e. The ratio between the average connection from *I* neurons and the one from *E* neurons was kept fixed with *w_I_/w_E_* = 5. We set the feedforward weight of input to I neurons to *w_I_*_f_ = 0.8. We simulated the dynamics of the model network using an Euler method with a time step of 0.5ms. Further details about the simulations and numerical estimates of mutual information and linear Fisher Information from simulated network responses can be found in Supplementary Information.

## Acknowledgements

National Institutes of Health grants 1R01MH115557 (K.J.), 1U19NS107613-01 (B.D.), R01EB026953 (B.D.); National Science Foundation grant NSF-DBI-1707400 (K.J.). Vannevar Bush faculty fellowship N00014-18-1-2002 (B.D); Simons Foundation Collaboration on the Global Brain (B.D.).

## Notes

### Competing Interest Statement

The authors have declared no competing interest.

