## Supplementary Information for "Recurrent circuit based neural population codes for stimulus representation and inference"

### 1 Suplementary Figures

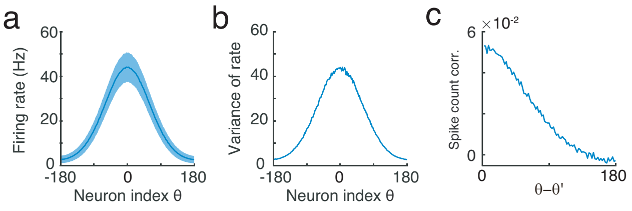

Figure S1: (a-b) The firing rate (a) and variance (b) of responses of E neurons in the network. The shaded area in (a) denotes the standard deviation. (c) The spike count correlation between two E neurons with their difference of preferred stimulus features.

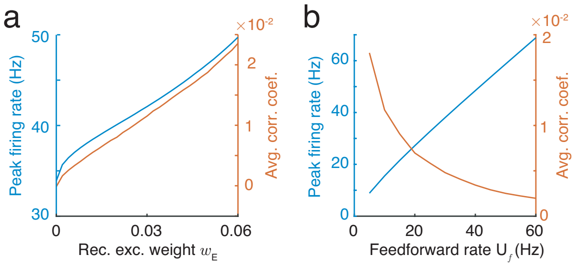

Figure S2: The peak firing rate (blue) and the averaged correlation coefficient (red) of excitatory neurons in the network model as a function of recurrent excitatory weight (a) and the feedforward input rate (b).

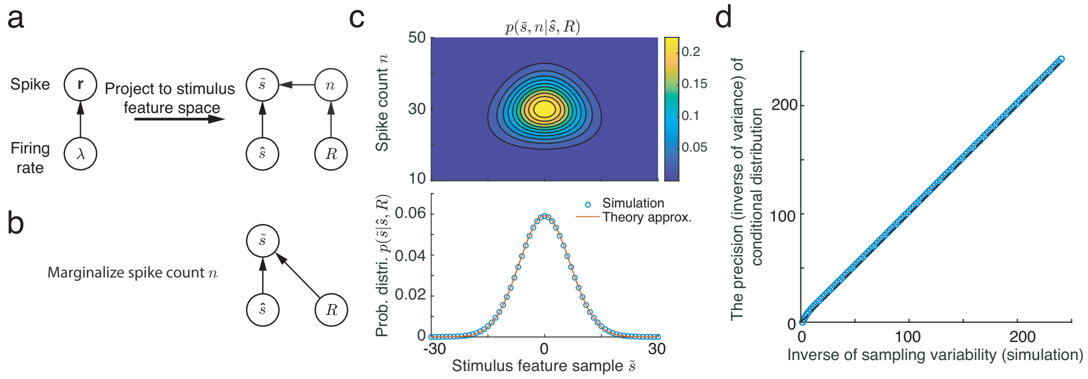

Figure S3: Poisson spiking variability provides correct sampling variability in stimulus feature space. (a) The likelihood function of stimulus feature embedded in the Poisson spike generation. The population firing rate $\boldsymbol{\lambda}$ is parameterized as a function over stimulus feature $s$ and peak firing rate $R$ (Eq. S16), and the stimulus sample $\tilde{s}$ and spike count $n$ are sufficient statistics from emitted spikes $\mathbf{r}$. (b) The trial averaged likelihood function of generating a sample of stimulus feature $\tilde{s}$. (c) The joint likelihood function $p(\tilde{s}, n|\hat{s}, R)$ (top) and the marginal likelihood $p(\tilde{s}|\hat{s}, R)$ (bottom). The theoretical approximation (red) is obtained by using Laplacian appximation (Eq. S21). (d) The variability of sampled stimulus feature averaged across trials is consistent with the variance of conditional distribution in Gibbs sampling.

#### 2 The equilibrium of Gibbs sampling dynamics

The iterations of Gibbs sampling algorithm can be analyzed theoretically by converting them into a discrete dynamics. We rewrite the Gibbs sampling dynamics (Eqs. 15-16 in maintext) at here for the convenience of readers.

$$
\begin{aligned}
p(s|\tilde{z}_t, \mathbf{u}^{\mathsf{f}}) &\propto \mathcal{N}\left(s|\hat{s}_t, \Lambda^{-1}\right), \qquad \text{(S1)}\\
\hat{s}_t &= \frac{\Lambda_{\mathsf{f}}\mu_{\mathsf{f}} + \Lambda_s \tilde{z}_t}{\Lambda_{\mathsf{f}} + \Lambda_s}, \quad \Lambda = \Lambda_{\mathsf{f}} + \Lambda_s. \\
\tilde{s}_t &\sim \mathcal{N}\left(s|\hat{s}_t, \Lambda^{-1}\right). \qquad \text{(S2)}\\
\tilde{z}_{t+\Delta t} &\sim \mathcal{N}(z|\tilde{s}_t, \Lambda_s^{-1}). \qquad \text{(S3)}
\end{aligned}
$$

For theoretical analysis, above three steps in the Gibbs sampling algorithm can be written as the iterations of the following discrete dynamical system,

$$
\begin{aligned}
\hat{s}_t &= \frac{\Lambda_{\mathsf{f}}\mu_{\mathsf{f}} + \Lambda_s \tilde{z}_t}{\Lambda_{\mathsf{f}} + \Lambda_s}, \\
\tilde{s}_t &= \hat{s}_t + (\Lambda_{\mathsf{f}} + \Lambda_s)^{-1/2}\xi_t, \\
\tilde{z}_{t+\Delta t} &= \tilde{s}_t + \Lambda_s^{-1/2}\epsilon_t,
\end{aligned} \qquad \text{(S4)}
$$

where $\epsilon_t$ and $\xi_t$ are independent samples from a standard normal distribution.

In Eq. (S4), the mean and variance of variables conditioned on $\mu_{\mathsf{f}}$ (the mean of the likelihood conveyed by feedforward input $\mathbf{u}^{\mathsf{f}}$, Eq. 11) in equilibrium state can be analytically calculated.

$$
\begin{aligned}
\langle \hat{s}|\mu_{\mathsf{f}}\rangle &= \mu_{\mathsf{f}}, \quad V(\hat{s}|\mu_{\mathsf{f}}) = \frac{\Lambda_s}{\Lambda_{\mathsf{f}}(\Lambda_{\mathsf{f}} + \Lambda_s)}, \\
\langle \tilde{s}|\mu_{\mathsf{f}}\rangle &= \mu_{\mathsf{f}}, \quad V(\tilde{s}|\mu_{\mathsf{f}}) = \Lambda_{\mathsf{f}}^{-1}, \\
\langle \tilde{z}|\mu_{\mathsf{f}}\rangle &= \mu_{\mathsf{f}}, \quad V(\tilde{z}|\mu_{\mathsf{f}}) = \Lambda_{\mathsf{f}}^{-1} + \Lambda_s^{-1}.
\end{aligned} \qquad \text{(S5)}
$$

#### 3 Theoretical analysis of neural population responses and Fisher information

##### 3.1 The conditional distribution of the neuronal response

We present the math details in computing the distribution of neuronal responses, $\mathbf{r}$, given an external stimulus feature $s$, i.e., $p(\mathbf{r}|s)$. For a fixed external stimulus, $s$, the neuronal response $\mathbf{r}$ fluctuates over time/trial due to both sensory transmission noise described by $p(\mathbf{u}^{\mathsf{f}}|s)$ (Eq. 13 and Fig. 2b), as well as the internally generated variability described by $p(\mathbf{r}|\mathbf{u}^{\mathsf{f}})$ (Fig. 3a). Hence we have,

$$
\begin{aligned}
p(\mathbf{r}|s) &= \int p(\mathbf{r}|\mathbf{u}^{\mathsf{f}})p(\mathbf{u}^{\mathsf{f}}|s)d\mathbf{u}^{\mathsf{f}}, \\
&= \int \Big[\int p(\mathbf{r}|\boldsymbol{\lambda})p(\boldsymbol{\lambda}|\mathbf{u}^{\mathsf{f}})d\boldsymbol{\lambda}\Big]p(\mathbf{u}^{\mathsf{f}}|s)d\mathbf{u}^{\mathsf{f}}. \qquad \text{(S6)}
\end{aligned}
$$

Furthermore, the internal variability $p(\mathbf{r}|\mathbf{u}^{\mathsf{f}})$ has two parts: One is $p(\mathbf{r}|\boldsymbol{\lambda})$ describing the independent Poisson spike generation (Eq. 20); and another is $p(\boldsymbol{\lambda}|\mathbf{u}^{\mathsf{f}})$ describing the fluctuation of instantaneous firing rate $\boldsymbol{\lambda}$ due to internal variability. We see that $p(\boldsymbol{\lambda}|\mathbf{u}^{\mathsf{f}})$ is the only unknown distribution in Eq. (S6), and we compute it in the text below.

Since we are mainly interested in how stimulus is represented in the stimulus feature subspace in neuronal response space, we only consider the covariability along the stimulus feature subspace, and ignore the covariability due to fluctuation along other directions. Therefore, the instataneous firing rate $\boldsymbol{\lambda}_t$ can be approximated as a smooth Gaussian profile over $\hat{s}_t$, where $\hat{s}_t$ is the mean of instantaneous conditional distribution (Eq. 4),

$$
\boldsymbol{\lambda}_{tj}(\hat{s}_t) \approx R\exp[-(\hat{s}_t-\theta_j)^2/(2a^2)].
$$

Hence the average of instantaneous firing rate $\boldsymbol{\lambda}_t$ over time corresponds to average of $\hat{s}_t$ over time,

$$
\begin{aligned}
\langle \boldsymbol{\lambda}_t(\hat{s}_t)|\mathbf{u}^{\mathsf{f}}\rangle &= \boldsymbol{\lambda}(\langle \hat{s}_t|\mathbf{u}^{\mathsf{f}}\rangle), \\
&= R\exp[-(\mu_{\mathsf{f}}-\theta_j)^2/(2a^2)] \triangleq \mathbf{f}(\mu_{\mathsf{f}}). \qquad \text{(S7)}
\end{aligned}
$$

The second equality in above equation comes from $\langle \hat{s}_t|\mathbf{u}^{\mathsf{f}}\rangle = \mu_{\mathsf{f}}$ as derived in Eq. (S5). Expanding the instantaneous firing rate $\boldsymbol{\lambda}_t(\hat{s})$ around the mean responses $\mathbf{f}(\mu_{\mathsf{f}})$,

$$
\boldsymbol{\lambda}(\hat{s}_t) = \mathbf{f}(\mu_{\mathsf{f}}) + \mathbf{f}'_{\mu_{\mathsf{f}}}(\hat{s}_t-\mu_{\mathsf{f}}),
$$

where $\mathbf{f}'_{\mu_{\mathsf{f}}} = d\mathbf{f}(\mu_{\mathsf{f}})/d\mu_{\mathsf{f}} = Ra^{-2}(\theta_j-\mu_{\mathsf{f}})\exp[-(\mu_{\mathsf{f}}-\theta_j)^2/(2a^2)]$, the covariance of instantaneous firing rate can be computed as

$$
\begin{aligned}
\Sigma[\boldsymbol{\lambda}(\hat{s})] &= \Big\langle [\boldsymbol{\lambda}(\hat{s}_t)-\mathbf{f}(\mu_{\mathsf{f}})][\boldsymbol{\lambda}(\hat{s}_t)-\mathbf{f}(\mu_{\mathsf{f}})]^\top \Big\rangle, \\
&= \big\langle (\hat{s}_t-\mu_{\mathsf{f}})^2 \big\rangle \mathbf{f}'_{\mu_{\mathsf{f}}}\mathbf{f}'^{\top}_{\mu_{\mathsf{f}}}, \\
&= V(\hat{s}|\mu_{\mathsf{f}})\mathbf{f}'_{\mu_{\mathsf{f}}}\mathbf{f}'^{\top}_{\mu_{\mathsf{f}}}. \qquad \text{(S8)}
\end{aligned}
$$

$V(\hat{s}|\mu_{\mathsf{f}})$ is the variance of $\hat{s}$ given the feedforward input $\mathbf{u}^{\mathsf{f}}$, as derived in Eq. (S5). The covariance structure $\mathbf{f}'_{\mu_{\mathsf{f}}}\mathbf{f}'^{\top}_{\mu_{\mathsf{f}}}$ captures the covariability due to firing rate fluctuations along the stimulus feature subspace, which is often termed differential correlations [1, 2]. With the Gaussian tuning of $\mathbf{f}(\mu_{\mathsf{f}})$

(Eq. S7), $\mathbf{f}'_{\mu_\mathsf{f}}\mathbf{f}'^{\top}_{\mu_\mathsf{f}}$ exhibits anti-symmetric structure over $\mu_\mathsf{f}$, i.e.,

$$\left[\mathbf{f}'_{\mu_\mathsf{f}}\mathbf{f}'^{\top}_{\mu_\mathsf{f}}\right]_{ij} \propto (\theta_i - \mu_\mathsf{f})(\theta_j - \mu_\mathsf{f}) \exp[-(\mu_\mathsf{f} - \theta_i)^2/(2a^2)] \exp[-(\mu_\mathsf{f} - \theta_j)^2/(2a^2)],$$

which is plotted in Fig. 6b. Combining Eqs. (S7 and S8) together, $p(\boldsymbol{\lambda}|\mathbf{u}^\mathsf{f})$ can be approximated as a multivariate normal distribution,

$$p(\boldsymbol{\lambda}|\mathbf{u}^\mathsf{f}) = \mathcal{N}\big[\boldsymbol{\lambda}\big|\mathbf{f}(\mu_\mathsf{f}), V(\hat{s}|\mu_\mathsf{f})\mathbf{f}'_{\mu_\mathsf{f}}\mathbf{f}'^{\top}_{\mu_\mathsf{f}}\big], \tag{S9}$$

where,

$$\begin{aligned} \mathbf{f}_j(\mu_\mathsf{f}) &= R\exp[-(\mu_\mathsf{f} - \theta_j)^2/2a^2], && \text{(S10)} \\ V(\hat{s}|\mu_\mathsf{f}) &= \frac{\Lambda_s}{\Lambda_\mathsf{f}(\Lambda_\mathsf{f} + \Lambda_s)} = a^2 n_\mathsf{f}^{-1} w_E^*. && \text{(S11)} \end{aligned}$$

Here we derive the Eq. (28) in the main text. Furthermore, by approximating the independent Poisson distributions $p(\mathbf{r}|\boldsymbol{\lambda})$ and $p(\mathbf{u}^\mathsf{f}|s)$ as multivariate normal distribution, i.e.,

$$p(\mathbf{r}|\boldsymbol{\lambda}) = \mathcal{N}[\mathbf{r}|\boldsymbol{\lambda}(\hat{s}), \mathrm{diag}(\boldsymbol{\lambda}(\hat{s}))],$$

the distribution $p(\boldsymbol{\lambda}|\mathbf{u}^\mathsf{f})$ can be computed as

$$\begin{aligned} p(\mathbf{r}|\mathbf{u}^\mathsf{f}) &= \int p(\mathbf{r}|\boldsymbol{\lambda})p(\boldsymbol{\lambda}|\mathbf{u}^\mathsf{f})d\boldsymbol{\lambda}, \\ &= \mathcal{N}\big[\mathbf{r}\big|\mathbf{f}(\mu_\mathsf{f}), \mathrm{diag}(\boldsymbol{\lambda}(\hat{s})) + V(\hat{s}|\mu_\mathsf{f})\mathbf{f}'_{\mu_\mathsf{f}}\mathbf{f}'^{\top}_{\mu_\mathsf{f}}\big]. && \text{(S12)} \end{aligned}$$

Compared with $p(\boldsymbol{\lambda}|\mathbf{u}^\mathsf{f})$ (Eq. S9), we see the covariance of $p(\mathbf{r}|\mathbf{u}^\mathsf{f})$ (Eq. S12) has an extra term, i.e., $\mathrm{diag}(\boldsymbol{\lambda}(\hat{s}))$, which is a diagonal matrix denoting the independent Poisson spiking variability. Finally, substituting Eqs. (S9) and (13) into Eq. (S6), shows that the response distribution conditioned on the external stimulus feature $s$, $p(\mathbf{r}|s)$, has the form

$$p(\mathbf{r}|s) \approx \mathcal{N}\big[\mathbf{r}\big|\mathbf{f}(s), \mathrm{diag}(\mathbf{f}(s)) + V(\hat{s}|s)\mathbf{f}'_s\mathbf{f}'^{\top}_s\big]. \tag{S13}$$

Here the variance $V(\hat{s}|s)$ in the stimulus feature subspace is a mixture of internal variability and sensory noise. It follows from Eqs. (S5 and 13) that

$$\begin{aligned} V(\hat{s}|s) &= V(\hat{s}|\mu_\mathsf{f}) + V(\mu_\mathsf{f}|s), \\ &= \frac{\Lambda_s}{\Lambda_\mathsf{f}(\Lambda_\mathsf{f} + \Lambda_s)} + \frac{1}{\Lambda_\mathsf{f}}, \\ &= a^2 n_\mathsf{f}^{-1}(w_E^* + 1). && \text{(S14)} \end{aligned}$$

The second term, $V(\mu_{\mathsf{f}}|s)$, represents fluctuations in neuronal activity in the stimulus feature subspace due to the sensory transmission noise (Eq. 13).

##### 3.2 Calclation of linear Fisher information in neuronal responses

We present the theoretical calculations of the amount of linear Fisher inforamtion of stimulus feature $s$ contained in neural population responses in a network implementing Gibbs sampling. From the neuronal response distribution (Eq. S13), the linear Fisher information of $s$ contained in $\mathbf{r}$ is [1],

$$\mathcal{I}(s|\mathbf{r}) \quad = \quad \mathbf{f}_s'^{\top}\Sigma(\mathbf{r}|s)^{-1}\mathbf{f}_s',$$

where $\mathbf{f}$ is the mean firing rate of neurons, and $\mathbf{f}_s'$ denotes the derivative of firing rate on the stimulus $s$. Using the matrix inverse lemma, the inverse of covariance matrix can be found as

$$\begin{aligned}\Sigma(\mathbf{r}|s)^{-1} &= \left[\mathrm{diag}(\mathbf{f}(s)) + V(\hat{s}|s)\mathbf{f}_s'^{\top}\mathbf{f}_s\right]^{-1}, \\ &= \mathrm{diag}(\mathbf{f}(s))^{-1} - V(\hat{s}|s)\frac{\mathrm{diag}(\mathbf{f}(s))^{-1}\mathbf{f}_s'\mathbf{f}_s'^{\top}\mathrm{diag}(\mathbf{f}(s))^{-1}}{1+V(\hat{s}|s)\mathbf{f}_s'^{\top}\mathrm{diag}(\mathbf{f}(s))^{-1}\mathbf{f}_s'}.\end{aligned}$$

Substituting $\Sigma(\mathbf{r}|s)^{-1}$ back into the calculation of $\mathcal{I}(s|\mathbf{r})$,

$$\begin{aligned}\mathcal{I}(s|\mathbf{r}) &= \mathbf{f}_s'^{\top}\mathrm{diag}(\mathbf{f}(s))^{-1}\mathbf{f}_s' - V(\hat{s}|s)\frac{(\mathbf{f}_s'^{\top}\mathrm{diag}(\mathbf{f}(s))^{-1}\mathbf{f}_s')^2}{1+V(\hat{s}|s)\mathbf{f}_s'^{\top}\mathrm{diag}(\mathbf{f}(s))^{-1}\mathbf{f}_s'}, \\ &= \frac{\mathbf{f}_s'^{\top}\mathrm{diag}(\mathbf{f}(s))^{-1}\mathbf{f}_s'}{1+V(\hat{s}|s)\mathbf{f}_s'^{\top}\mathrm{diag}(\mathbf{f}(s))^{-1}\mathbf{f}_s'}.\end{aligned}$$

Substituting the result of $V(\hat{s}|s)$ (Eq. S14) and Eq. (24) into above equation,

$$\mathcal{I}(s|\mathbf{r}) = \frac{\Lambda_{\mathsf{f}}}{2}.$$

##### 3.3 Invertible linear transformations do not degrade linear Fisher information

A network that linearly transforms its inputs does not degrade linear Fisher information. On the other hand, internally generated differential correlations will decrease linear Fisher information. To prove this, we write the activity of cells in a linear network in response to a stimulus $s$ as,

$$\mathbf{r}(s) = \mathbf{A}\mathbf{u}^{\mathsf{f}}(s) + \epsilon\boldsymbol{\eta}.$$

Here $\mathbf{A}$ is the linear transformation matrix and is determined by the recurrent connections in the network, $\mathbf{u}^{\mathsf{f}}(s)$ is the feedforward input defined in Eq. (10). The term $\epsilon\boldsymbol{\eta}$, captures the internally generated differential correlations with amplitude controlled by $\epsilon$ (Eq. 7). The random variable $\boldsymbol{\eta}$ has zero mean, and covariance equal to $\Sigma_{\boldsymbol{\eta}} = \mathbf{f}_s'\mathbf{f}_s'^{\top}$, with $\mathbf{f}_s' = d\mathbf{f}(s)/ds$, and $\mathbf{f}(s) = \langle\mathbf{r}(s)\rangle$ is the tuning of network responses (Eq. 7). When the matrix $\mathbf{A}$ is *invertible*, the linear Fisher information of $s$ in neuronal response $\mathbf{r}$ is,

$$\begin{aligned}\mathcal{I}_{\mathbf{r}}(s) &= \mathbf{f}_s'^{\top}\Sigma_{\mathbf{r}}^{-1}\mathbf{f}_s',\\ &= \left(\mathbf{A}\langle\mathbf{u}^{\mathsf{f}}\rangle_s'\right)^{\top}\left[\mathbf{A}\Sigma_{\mathbf{u}^{\mathsf{f}}}\mathbf{A}^{\top}+\epsilon\mathbf{f}_s'\mathbf{f}_s'^{\top}\right]^{-1}\mathbf{A}\langle\mathbf{u}^{\mathsf{f}}\rangle_s',\\ &= \langle\mathbf{u}^{\mathsf{f}}\rangle_s'^{\top}\left[\Sigma_{\mathbf{u}^{\mathsf{f}}}+\epsilon\langle\mathbf{u}^{\mathsf{f}}\rangle_s'\langle\mathbf{u}^{\mathsf{f}}\rangle_s'^{\top}\right]^{-1}\langle\mathbf{u}^{\mathsf{f}}\rangle_s',\end{aligned}$$

where $\langle\mathbf{u}^{\mathsf{f}}\rangle$ is the tuning of feedforward inputs, $\mathbf{u}^{\mathsf{f}}$ (Eq. 10).

Thus the linear Fisher information does not depend on $\mathbf{A}$, and hence a linear invertible transformation does not decrease the linear Fisher information. By using the matrix inverse lemma we find that

$$\left[\Sigma_{\mathbf{u}^{\mathsf{f}}}+\epsilon\langle\mathbf{u}^{\mathsf{f}}\rangle_s'\langle\mathbf{u}^{\mathsf{f}}\rangle_s'^{\top}\right]^{-1}=\Sigma_{\mathbf{u}^{\mathsf{f}}}^{-1}-\frac{\epsilon}{1+\epsilon\langle\mathbf{u}^{\mathsf{f}}\rangle_s'^{\top}\Sigma_{\mathbf{u}^{\mathsf{f}}}^{-1}\langle\mathbf{u}^{\mathsf{f}}\rangle_s'}\Sigma_{\mathbf{u}^{\mathsf{f}}}^{-1}\langle\mathbf{u}^{\mathsf{f}}\rangle_s'\langle\mathbf{u}^{\mathsf{f}}\rangle_s'^{\top}\Sigma_{\mathbf{u}^{\mathsf{f}}}^{-1},$$

the linear Fisher information of stimulus $s$ in $\mathbf{r}$ can be computed as,

$$\mathcal{I}_{\mathbf{r}}(s)=\frac{\langle\mathbf{u}^{\mathsf{f}}\rangle_s'^{\top}\Sigma_{\mathbf{u}^{\mathsf{f}}}^{-1}\langle\mathbf{u}^{\mathsf{f}}\rangle_s',}{1+\epsilon\langle\mathbf{u}^{\mathsf{f}}\rangle_s'^{\top}\Sigma_{\mathbf{u}^{\mathsf{f}}}^{-1}\langle\mathbf{u}^{\mathsf{f}}\rangle_s'}=\frac{\mathcal{I}_{\mathbf{u}^{\mathsf{f}}}(s)}{1+\epsilon\mathcal{I}_{\mathbf{u}^{\mathsf{f}}}(s)}. \tag{S15}$$

If the linear network doesn't internally generate differential correlation, i.e., $\epsilon=0$, the linear Fisher information in the network response is the same as the the information inherited from feedforward input, i.e., $\mathcal{I}_{\mathbf{r}}(s)=\mathcal{I}_{\mathbf{u}^{\mathsf{f}}}(s)$. Hence internal differential correlations degrade linear Fisher information, but any linear invertible transformations by the network do not.

### 4 Poisson spiking variability correctly samples in stimulus feature subspace

We present the math details in demonstrating the Poisson spiking variability provides correct variability for sampling stimulus feature when averaged over trials. That is, the variability in the stimulus feature subspace from the Poisson spiking variability is consistant with the variance of conditional distribution (Eq. 16). To simplify the math proof, we consider the instantaneous firing rate $\boldsymbol{\lambda}_t$ has a smooth Gaussian profile over $\hat{s}_t$, the mean of the instantaneous conditional distribution of $s$ (Eq. 16), i.e.,

$$\boldsymbol{\lambda}_{tj}(\hat{s}_t,R)\approx R\exp[-(\hat{s}_t-\theta_j)^2/2a^2], \tag{S16}$$

and $\boldsymbol{\lambda}_{tj}$ can be effectively treated as a deterministic function over $\hat{s}_t$ and $R$. This smooth Gaussian approximation can be satisfied in a sufficient long time window or there is large number of neurons having the same preference over stimulus feature $\theta$. Substituting Eq. (S16) into Eq. (20), the likelihood of $\hat{s}$ and $R$ can be derived as (the time index $t$ is omitted to simplify the notation),

$$p(\mathbf{r}|\boldsymbol{\lambda}) \propto p(\mathbf{r}|\hat{s}, R) \propto R^{\sum_j r_j} \exp\Big[-\sum_j r_j \tfrac{(\hat{s}-\theta_j)^2}{2a^2}\Big] \exp\Big[-R\sum_j e^{-(\hat{s}-\theta_j)^2/2a^2}\Big]. \tag{S17}$$

To simplify notations, we denote

$$n = \sum_j \mathbf{r}_j, \quad \beta = \sum_j e^{-(\hat{s}-\theta_j)^2/2a^2},$$

where $n$ is the spike count of emitted spikes, and $\beta$ is the sum of normalized firing rate of all neurons. With uniform distribution of the preference of neurons $\{\theta_j\}_{j=1}^N$ in stimulus feature subspace as assumed in main text, $\beta$ will be a constant value irrelevant with the amount of input spikes. Normalizing the distribution in Eq. (S17), the likelihood of $\hat{s}$ and $R$ can be derived as (Fig. S2A),

$$\begin{aligned} p(\tilde{s}, n|\hat{s}, R) &\propto \mathcal{N}\left(\hat{s}|\tilde{s}, a^2 n^{-1}\right) \Gamma(R|n+1, \beta), \\ &\propto \mathcal{N}\left(\tilde{s}|\hat{s}, a^2 n^{-1}\right) \text{Poisson}(n|\beta R), \end{aligned} \tag{S18}$$

where $\Gamma(\cdot)$ denotes a Gamma distribution in above equation. $\tilde{s}$ can be regarded as a sample of stimulus feature, which is can be read out from the generated Poisson spikes $\mathbf{r}$,

$$\tilde{s} = n^{-1} \sum_j \mathbf{r}_j \theta_j.$$

Eq. (S18) suggests the variability of generating a sample of stimulus feature $\tilde{s}$ conditioned on $\hat{s}$ is $a^2 n^{-1}$ which is inversely proportional to the number of emitted spikes $n$. In comparison, the variance of conditional distribution is $\Lambda^{-1} = a^2(\sum_j \lambda_j)^{-1}$, which is inversely proportional to the sum of instantaneous firing rate (Eq. 4 and Table 2). Therefore, determining whether Poisson spike variability correctly samples a stimulus feature is equivalent to compare whether the emitted spike count $n$ is equal to the sum of firing rate $\sum_j \lambda_j$. Although in a single trial $n$ is not necessarily the same as $\sum_j \lambda_j$, we notice that,

$$\langle n \rangle = \beta R = \sum_j \boldsymbol{\lambda}_j,$$

i.e., the trial average of spike count $n$ when fixing the instantaneous firing rate $\boldsymbol{\lambda}$ is the same as $\boldsymbol{\lambda}$ (Eq. S18). This suggests the variability from Poisson spike generation averaged over trials is consistent with the variance of conditional distribution.

To rigorously prove the Poisson spiking variability averaged over trials correctly samples a stimulus feature, we compute the trial averaged likelihood of generating a stimulus sample $\tilde{s}$ given an instantaneous firing rate $\boldsymbol{\lambda}_t$ (corresponding to a given $\hat{s}$ and $R$). This corresponds to marginalize the number of emitted spike count $n$,

$$
\begin{aligned}
p(\tilde{s}|\boldsymbol{\lambda}) &= \sum_n p(\tilde{s}, n|s, R), \\
&= \sum_n \mathcal{N}(\tilde{s}|\hat{s}, a^2 n^{-1}) \mathrm{Poisson}(n|\beta R).
\end{aligned}
$$

To simplify the analytical calculation, we approximate the Poisson distribution in above equation by a continuous Gaussian distribution, which works well when the summed firing rate over all neurons $\beta R$ is not so small.

$$
\begin{aligned}
p(\tilde{s}|\hat{s}, R) &\approx \int \mathcal{N}\left(\tilde{s}|\hat{s}, a^2 n^{-1}\right) \mathcal{N}(n|\beta R, \beta R) dn, \\
&\approx \mathcal{N}\left(\tilde{s}|\hat{s}, a^2 \hat{n}^{-1}\right) \mathcal{N}(\hat{n}|\beta R, \beta R) \det(\mathrm{H}/2\pi)^{-1/2},
\end{aligned} \tag{S19}
$$

We use the Laplace's methods to approximate the integral in above equation [3], which yields the approximation on the second row in above equation, where

$$
\hat{n} = \arg\max_n p(\tilde{s}, n|\hat{s}, R), \quad \mathrm{H} = -\frac{\partial^2 \ln p(\tilde{s}, n|\hat{s}, R)}{\partial n^2}\Big|_{n=\hat{n}}. \tag{S20}
$$

After some tedious analytical calculation (see Appendix 7.1), the trial averaged likelihood of a sample of stimulus feature can be approximated as a Gaussian distribution (Fig. S3b),

$$
p(\tilde{s}|\hat{s}, R) \approx \mathcal{N}\left[\tilde{s}\middle|\hat{s}, a^2 (\textstyle\sum_j \boldsymbol{\lambda}_j)^{-1}\right], \tag{S21}
$$

whose variance is the same as the variance of conditional distribution (Eq. 4). Therefore, the Poisson spiking variability provides correct variability for sampling stimulus feature asymptotically. Fig. S3c-d verifies that the Laplacian approximation used in theoretical analysis works very well as long as the peak firing rate $R$ is not too small.

### 5 Hybrid representation in the network

The proposed network has a hybrid mechanism in representing probability distributions, in which the posterior of latent hierarchical features is approximated by sequential samples, and the instantaneous conditional distribution of stimulus features is parametrically represented by instantaneous firing rate.

#### 5.1 Posterior is approximated by sequential samples

Since the proposed network (Eqs. 19-20) embeds the Gibbs sampling into its dynamics, the marginal posterior of stimulus feature is approximated by the sequential samples, and each sample can be read out from instantaneous spiking response $\mathbf{r}_t$ via a linear decoder,

$$p(s|x) \approx \frac{1}{T}\sum_t \delta(s - \tilde{s}_t), \quad \tilde{s}_t = \sum_j \mathbf{r}_{tj}\theta_j / \sum_j \mathbf{r}_{tj},$$

where $T$ is the number of samples, and $\delta(\cdot)$ is the Dirac delta function. Similarly, the marginal posterior of context feature is also approximated by the sequential samples as

$$p(z|x) \approx \frac{1}{T}\sum_t \delta(z - \tilde{z}_t), \quad \tilde{z}_t = \sum_j \mathbf{u}_{r,tj}\theta_j / \sum_j \mathbf{u}_{r,tj},$$

If we are interested in the joint posterior, which could be approximated by combining the samples of stimulus and context features together,

$$p(s,z|x) \approx \frac{1}{T}\sum_t \delta\left[(s,z)^\top - (\tilde{s}_t, \tilde{z}_t)^\top\right]$$

Combined together, the posterior distribution is represented by the sequantial samples in the proposed network.

#### 5.2 Instantaneous firing rate parametrically represents the conditional distribution

In the proposed network model, Eq. 4 suggets that the instantaneous firing rate $\boldsymbol{\lambda}_t$ parametrically represents the instantaneous conditional distribution $p(s|\tilde{z}_t, \mathbf{u}^\mathrm{f})$ via a log-probability code, in which the logarithm of $p(s|\tilde{z}_t, \mathbf{u}^\mathrm{f})$ is a linear function of the instataneous firing rate $\boldsymbol{\lambda}_t$ (Eq. 4), i.e.,

$$\ln p(s|\tilde{z}_t, \mathbf{u}^\mathrm{f}) = \ln \mathcal{N}(s|\hat{s}_t, \Lambda^{-1}) = -\sum_j \boldsymbol{\lambda}_{tj} \frac{(s-\theta_j)^2}{2a^2}. \tag{S22}$$

Hence $\boldsymbol{\lambda}_t$ contains all the information of the instantaneous conditional distribution.

#### 5.3 Mean response: mean-field approximation of the conditional distribution

The instantaneous firing rate $\boldsymbol{\lambda}_t$ is a hypothetical variable in a Poisson neuron model, but it cannot be directly observed. In neural data analysis, it is usual to estimate the mean neuronal responses (spike count) averaged over time or over trial, i.e., $\langle \mathbf{r}_t \rangle$, which is equal to the averaged instantaneous firing rate, i.e.,

$$\langle \mathbf{r}_t \rangle = \langle \boldsymbol{\lambda}_t \rangle.$$

In the proposed network, the mean responses $\langle \mathbf{r}_t \rangle$ parametrically represents the time averaged conditional distribution. By averaging Eq. (S22), using the relation in above equation, we have

$$
\begin{aligned}
&- \sum_j \langle \boldsymbol{\lambda}_t \rangle \frac{(s-\theta_j)^2}{2a^2} = \langle \ln p(s|\hat{s}_t, \Lambda) \rangle_t \,, \\
\Leftrightarrow &- \sum_j \langle \mathbf{r}_t \rangle \frac{(s-\theta_j)^2}{2a^2} \approx \ln p(s|\langle \hat{s}_t \rangle, \Lambda).
\end{aligned} \tag{S23}
$$

A mean-field approximation is used in the second row in Eq. (S23), in which the expectation of the log distribution is approximated by a new log distribution with the expectation of parameter [3]. This approximation works well when the fluctuation of $\tilde{s}_t$ over time is much smaller than the variance of the distribution, i.e., $V(\tilde{s}_t) \ll \Lambda^{-1}$.

The mean field-approximation suggests the time averaged conditional distribution of stimulus feature, i.e., $p(s|\langle \hat{s}_t \rangle, \Lambda)$, could be read out from the averaged neuronal responses. From Eq. (S23), the time averaged conditional distribution could be read out as,

$$
\langle \tilde{s}_t \rangle = \frac{\sum_j \langle \mathbf{r}_{tj} \rangle_t \theta_j}{\sum_j \langle \mathbf{r}_{tj} \rangle_t}, \quad \Lambda = a^{-2} \sum_j \langle \mathbf{r}_{tj} \rangle.
$$

### 6 Estimating the information content in network simulation

To estimate the information context contained in network responses, we fix an external stimulus feature $s$ in the world, we generated independent Poissonian feedforward inputs on every time step, and simulated the network dynamics accordingly (Eqs. 31-34). For simplicity, we set the external stimulus feature $s$ to zero without loss of generality, since the feedforward inputs and network responses are translation-invariant.

In estimating the mutual information (Eq. 17) in network responses, we simulate the spiking network (Eqs. 31-34) for 2000 seconds. On each time step, the samples of the stimulus feature and context feature were read out by a population vector from the instantaneous spiking activities of E neurons, $\mathbf{r}^E$, and recurrent input of E neurons, $\mathbf{u}^{E\mathsf{r}}$ respectively (Table 2),

$$
\tilde{s}_t = \textstyle\sum_j \mathbf{r}_j^E(t)\theta_j / \sum_j \mathbf{r}_j^E(t), \quad \tilde{z}_t = \sum_j \mathbf{u}_j^{E\mathsf{r}}(t)\theta_j / \sum_j \mathbf{u}_j^{E\mathsf{r}}(t).
$$

Then the joint distribution of samples $\tilde{s}$ and $\tilde{z}$ is approximated as a bivariate normal distribution, and its mean, $\boldsymbol{\mu}_q$, and covariance matrix, $\mathbf{K}_q$, are computed. Note that the posterior (Eq. 14) is defined on a given feedforward input $\mathbf{u}^{\mathsf{f}}$ rather than a given external stimulus feature $s$. The fluctuation of $\mathbf{u}^{\mathsf{f}}$ over time will bring extra variability in the distribution of samples generated by the network compared with posterior. If a network optimally samples posterior, the mean and precision matrix of samples read out from the network in response to a fixed external stimulus feature $s$ are,

$$\boldsymbol{\mu}_q = (\langle \mu_{\mathsf{f}} \rangle, \langle \mu_{\mathsf{f}} \rangle)^\top = (s, s)^\top, \quad \mathbf{K}_q = \begin{pmatrix} \frac{1}{2}\Lambda_{\mathsf{f}} + \Lambda_s & -\Lambda_s \\ -\Lambda_s & \Lambda_s \end{pmatrix}.$$

Comparing these two equations with Eq. (14), we see the temporal fluctuations in $\mathbf{u}^{\mathbf{f}}$ only change the first element in precision matrix $\mathbf{K}_q$. To cancel its effect, we add the first element of $\mathbf{K}_q$ with $\Lambda_{\mathsf{f}}/2$, where $\Lambda_{\mathsf{f}}$ is read out from feedforward inputs (Eq. 12). Then the mean and corrected precision matrix of samples are substituted into Eq. (18) to get the mutual information contained in the population activities about $\Theta = \{s, z\}$.

The linear Fisher information of stimulus feature $s$ in responses of E neurons, $\mathbf{r}^E$, is estimated by using an bias-correction algorithm [4]. A pair of stimuli $s_\pm = \pm 1^\circ$ are individually presented to the network for $T = 1 \times 10^4$ trials with each trial lasting 200ms, and the empirical mean and covariance of E neurons' responses $\mathbf{r}^E$ under each stimulus are recorded. Then the bias-correction estimate of linear Fisher information is,

$$\mathcal{I}_{bc}(s) = \frac{d\langle \mathbf{r}^E \rangle}{ds}^\top \Sigma(\mathbf{r}^E)^{-1} \frac{d\langle \mathbf{r}^E \rangle}{ds} \frac{2T - N - 3}{2T - 2} - \frac{2N}{T ds^2}, \tag{S24}$$

where $d\langle \mathbf{r} \rangle = \langle \mathbf{r}_+^E \rangle - \langle \mathbf{r}_-^E \rangle$, $ds = s_+ - s_-$, and $\langle \cdot \rangle$ represents the average over trials. $\Sigma(\mathbf{r}^E)$ is the covariance matrix of E neurons' responses. $N$ is the number of neurons in the network.

### 7 Appendix

#### 7.1 Computing the averaged likelihood by using Laplace's method

We present the math details in computing the $\hat{n}$ and H defined in Eq. (S20) in order to approximate the integral in Eq. (S19) by using Laplace's method [3]. To simplify notations, we denote $\mathcal{L} \equiv p(\tilde{s}, n|\hat{s}, R)$, and then,

$$\ln \mathcal{L} \quad = \quad \frac{1}{2}\ln n - \frac{n}{2a^2}(\tilde{s}-\hat{s})^2 - \frac{(n-\beta R)^2}{2\beta R}. \tag{A1}$$

$\hat{n}$ can be found by taking the derivative of $\mathcal{L}$ over $n$ to be zero,

$$\frac{\partial \ln \mathcal{L}}{\partial n} \quad = \quad \frac{1}{2n} - \frac{(\tilde{s}-\hat{s})^2}{2a^2} - \frac{n-\beta R}{\beta R} = 0,$$

then the $\hat{n}$ can be computed as

$$\begin{aligned} \hat{n} \quad &= \quad \left[1-\frac{(\tilde{s}-\hat{s})^2}{2a^2}\right]\frac{\beta R}{2} + \frac{1}{2}\sqrt{\left[1-\frac{(\tilde{s}-\hat{s})^2}{2a^2}\right]^2\beta^2 R^2 + 2\beta R}, \\ &\approx \quad \left[1-\frac{(\tilde{s}-\hat{s})^2}{2a^2}\right]\beta R + \mathcal{O}(R^{1/2}). \end{aligned} \tag{A2}$$

To gain theoretical insight, we simplify above equation by omitting the term $2\beta R$ inside the square root function . This approximation works well when $R$ is large enough, since $R^2$ is larger than $R$ by an order. Meanwhile, the negative Hessian matrix is,

$$\begin{aligned} \frac{\partial^2 \mathcal{L}}{\partial n^2}\Big|_{n=\hat{n}} \quad &= \quad -\frac{1}{2\hat{n}^2} - \frac{1}{R}, \\ &= \quad -\frac{1}{2[1-\frac{(\tilde{s}-\hat{s})^2}{2a^2}]^2\beta^2 R^2} - \frac{1}{R}, \\ &\approx \quad -\frac{1}{R} + \mathcal{O}(R^{-2}). \end{aligned} \tag{A3}$$

This approximation is also considered under the large $R$ limit where the omitted term is a order smaller than $1/R$.

Substituting Eqs. (A2 and A3) back into Eq. (S19), we get an unnormalized likelihood for moving direction $x$,

$$p(\tilde{s}|\hat{s}, R) \propto \sqrt{\hat{n}(\tilde{s})}\exp\left[-\frac{\hat{n}(\tilde{s})}{2a^2}(\tilde{s}-\hat{s})^2\right]\exp\left[-\frac{(\hat{n}(\tilde{s})-\beta R)^2}{2\beta R}\right]. \tag{A4}$$

Since this distribution is complicated, again, we approximate it by a Gaussian distribution. It is easy to see $p(\tilde{s}|\hat{s}, R)$ is a symmetric distribution over its center $\tilde{s} = \hat{s}$, and it is a mixture of Gaussian distributions with different width (Fig. S3c). Next, we compute the Hessian of $p(\tilde{s}|\hat{s}, R)$ at it peak location, i.e., $\tilde{s} = \hat{s}$, which tells of the precision fo the Gaussian distribution approximating $p(\tilde{s}|\hat{s}, R)$. Denote by $\mathcal{L}(\tilde{s}) = p(\tilde{s}|\hat{s}, R)$ to simplify notations, and we have,

$$\ln \mathcal{L}(\tilde{s}) = \frac{1}{2} \ln \hat{n}(\tilde{s}) - \frac{\hat{n}(\tilde{s})}{2a^2}(\tilde{s} - \hat{s})^2 - \frac{(\hat{n}(\tilde{s}) - \beta R)^2}{2\beta R}. \tag{A5}$$

The Hessian of $\mathcal{L}(\tilde{s})$ is calculated as

$$\begin{aligned}
\frac{\partial \ln \mathcal{L}(\tilde{s})}{\partial \tilde{s}} &= \frac{1}{2\hat{n}(\tilde{s})}\frac{\partial \hat{n}(\tilde{s})}{\partial \tilde{s}} - \frac{(\tilde{s}-\hat{s})^2}{2a^2}\frac{\partial \hat{n}(\tilde{s})}{\partial \tilde{s}} - \frac{\hat{n}(\tilde{s})}{a^2}(\tilde{s}-\hat{s}) - \frac{\hat{n}(\tilde{s}) - \beta R}{\beta R}\frac{\partial \hat{n}(\tilde{s})}{\partial \tilde{s}}, \\
\frac{\partial^2 \ln \mathcal{L}(\tilde{s})}{\partial \tilde{s}^2} &= -\left(\frac{1}{2\hat{n}(\tilde{s})^2} + \frac{1}{\beta R}\right)\left(\frac{\partial \hat{n}(\tilde{s})}{\partial \tilde{s}}\right)^2 + \left[\frac{1}{2\hat{n}(\tilde{s})} - \frac{(\tilde{s}-\hat{s})^2}{2a^2} - \frac{\hat{n}(\tilde{s})}{\beta R} + 1\right]\frac{\partial^2 \hat{n}(\tilde{s})}{\partial \tilde{s}^2} \\
&\quad -2\frac{(\tilde{s}-\hat{s})}{a^2}\frac{\partial \hat{n}(\tilde{s})}{\partial \tilde{s}} - \frac{1}{a^2}\hat{n}(\tilde{s}).
\end{aligned} \tag{A6}$$

Meanwhile, the derivative of $\hat{n}(\tilde{s})$ over $\tilde{s}$ is

$$\frac{\partial \hat{n}(\tilde{s})}{\partial \tilde{s}} = \frac{\beta R}{a^2}(\hat{s} - \tilde{s}), \quad \frac{\partial^2 \hat{n}(\tilde{s})}{\partial \tilde{s}^2} = -\frac{\beta R}{a^2}. \tag{A7}$$

Substituting Eq. (A7) into Eq. (A6), the the Hessian of $\mathcal{L}(\tilde{s})$ at $\tilde{s} = \hat{s}$ can be calculated as,

$$\left.\frac{\partial^2 \ln \mathcal{L}(\tilde{s})}{\partial \tilde{s}^2}\right|_{\tilde{s}=\hat{s}} = -a^{-2}\left(\frac{1}{2} + \beta R\right) \approx -a^{-2}\beta R. \tag{A8}$$

We throw out 1/2 inside the parenthesis in above equation since it is much smaller than $\beta R$ when $R$ is large. Finally, the likelihood can be approximated as a Gaussian distribution

$$p(\tilde{s}|\hat{s}, R) \;\approx\; \mathcal{N}\left[\tilde{s}|\hat{s}, a^2(\beta R)^{-1}\right], \tag{A9}$$

which turns to be Eq. (S21). Note that $\beta R = \sum_j \mathbf{f}_j$ is the sum of firing rate, which suggets the variance of the trial averaged likelihood is consistent with the variance of conditional distribution.
